# Phenotypic and Transcriptomic Characterization of ElyC-Defective *Escherichia coli* Cells Reveal the Importance of ElyC in Cell Envelope Biology at Optimal and Sub-Optimal Temperatures

**DOI:** 10.1101/2024.04.10.588480

**Authors:** Fardin Ghobakhlou, Imène Kouidmi, Laura Alvarez, Felipe Cava, Catherine Paradis-Bleau

## Abstract

The bacterial cell envelope acts as the frontline defense against environmental and internal stress, maintaining cellular homeostasis. Understanding envelope biology is crucial for both fundamental research and practical applications. Peptidoglycan (PG) is a key structural element, protecting against mechanical and osmotic stress while maintaining cell shape and integrity. In a previous study, we discovered the importance of ElyC, a highly conserved *Escherichia coli* protein with an unknown function, in maintaining envelope integrity at low temperatures. ElyC is essential for PG assembly at 21°C and plays a role in lipid carrier metabolism, a crucial step in PG and other bacterial envelope polysaccharide biosynthesis. At 21°C, ElyC deficiency leads to complete PG assembly blockage and cell lysis. However, the significance of ElyC in cells grown at 37°C remained unexplored. In our recent study, we conducted phenotypic and transcriptomic profiling of ElyC-defective *E. coli* cells grown at 37°C and 21°C, compared to wild-type cells. While *ΔelyC* mutant cells grow similarly to wild-type cells at 37°C, microscopy revealed altered cell morphology due to ElyC’s absence. PG quantification confirmed significantly inhibited PG biosynthesis at 37°C without ElyC, and these mutants showed increased sensitivity to PG-targeting β-lactam antibiotics compared to wild-type cells at the same temperature. RNA-Seq analysis of *ΔelyC* mutant and WT strains at 21°C and 37°C revealed that ElyC deletion severely affects the cell envelope at 21°C and moderately at 37°C. Several pathways and genes, especially stress response pathways, impact cell envelope functions, including biogenesis, maintenance, repair, metabolism, respiratory chain, peptidoglycan, lipopolysaccharide, membrane, cell wall, oxidative stress, osmotic stress, trehalose, chaperone, oxidoreductase, amino sugar synthesis and metabolism, vancomycin and beta-lactam resistance pathways and are affected. Downregulated transcripts are associated with mobility, arginine metabolism, membrane transport, regulation, outer membrane, transferase, and unknown functions. Our data highlights ElyC’s broad role in bacterial cell envelope and peptidoglycan biosynthesis at varying temperatures.

**IMPORTANCE:** The molecular pathways governing bacterial envelope biosynthesis, assembly, regulation, and adaptation remain incompletely understood. Envelope biology is vital for both fundamental microbiological research and the development of novel therapeutic targets. We previously established ElyC’s role in sub-optimal temperature envelope biology, showing its essentiality for PG assembly and bacterial survival at 21°C. In this study, we show that ElyC, a protein containing the highly conserved DUF218 domain of unknown function, is crucial for proper cell morphology, PG biosynthesis, antibiotic tolerance and envelope homeostasis at 37°C. Our findings emphasize the significance of DUF218-containing ElyC in envelope biology at physiological temperatures and uncover a novel cold-sensitive process in bacterial envelope biology.

## INTRODUCTION

The bacterial cell envelope facilitates selective nutrient influx and toxin efflux, supporting both catabolic and anabolic processes. This functional structure acts as the frontier line of defense, mediating signal transduction and cellular adaptation in response to environmental and intrinsic stressors to maintain cellular homeostasis. Additionally, it orchestrates morphogenesis, cell division, energy production, mobility, pathogenesis, and antibiotic resistance. The envelope of Gram-negative bacteria is made of an inner and an outer membrane that delimits the periplasmic space containing a thin layer of peptidoglycan (PG). Each layer of the cell envelope processes its own unique composition, organization, and functions, making it a remarkable biological structure {Mitchell, 2019 #51; Dufresne, 2015 #3}.

The inner membrane (IM) consists of a symmetric phospholipid bilayer with helical-type proteins and lipoproteins, playing several essential physiological roles. Among these functions, it drives energy production through the proton gradient created via the electron transfer chain, enabling the production of the proton motive force in the IM and the ATP in the cytoplasm {Dufresne, 2015 #3}. Another key component of the Gram-negative bacterial envelope is peptidoglycan, a unique and essential element. It encases the inner membrane and provides partial protection against cell rupture due to internal turgor and mechanical load {Rojas, 2018 #63}. The primary structural components of the peptidoglycan are long glycan strands composed of alternating units of N-acetylglucosamine (GlcNAc) and N-acetylmuramic acid (MurNAc), cross-linked with short peptides {Vollmer, 2008 #6}. Although the peptidoglycan is virtually universal in bacteria, its chemical structure exhibits diversity {Yadav, 2018 #71}. Recent molecular imaging studies using atomic force microscopy (AFM) combined with size exclusion chromatography have shown that the orientation of glycan chains in peptidoglycan is less ordered than previously described. Also, alteration in cell shape is associated with changes in the biophysical properties of peptidoglycan. When *E. coli* is in its normal rod shape, glycans are long and oriented, but when a spheroid shape is induced (chemically or genetically) glycans become short and disordered {Turner, 2018 #68}. The modifications in both sugars and peptides are mechanisms by which bacteria to cope with diverse environmental challenges. The peptidoglycan biosynthesis process occurs in different cell compartments. The biosynthesis of the final lipid-linked precursor of peptidoglycan (lipid II) begins in the cytoplasm and is completed in the inner leaflet of the inner membrane. The lipid II consists of β1,4-linked GlcNAc and MurNAc, with MurNAc linked to both the pentapeptide and the undecaprenyl phosphate lipid carrier (Und-P) {Vollmer, 2008 #7}. The lipid II is subsequently translocated to the outer leaflet of the inner membrane by the flippase MurJ {Meeske, 2015 #8} and integrated into the growing PG network by the high molecular weight Penicillin-Binding Proteins (PBPs) {Sauvage, 2008 #9, which are the targets of penicillin and other β-lactam related antibiotics. The outer membrane (OM) is an asymmetrical lipid bilayer with phospholipids in the inner leaflet and mainly lipopolysaccharides (LPS_s_) in the outer leaflet {Abellon-Ruiz, 2017 #124;Shearer, 2019 #125}. The Mla pathway, a six-component system found in many Gram-negative bacteria, maintains this asymmetry by transporting misplaced phospholipids from OM to the inner membrane (IM) {Abellon-Ruiz, 2017 #124}. Furthermore, the Mla system, along with an ABC transporter of unknown function (YadH), preserves outer membrane lipid asymmetry {Babu, 2018 #66}. The OM contains various proteins known as outer membrane proteins (OMPs), which serve essential functions for the cell, including nutrient uptake, cell adhesion, cell signaling, waste export, acting as virulence factors, and providing a protective barrier {Mitchell, 2019 #74} {Dufresne, 2015 #51}{Rollauer, 2015 #126}. A research has shown that the cell wall is not the dominant mechanical element within Gram-negative bacteria, instead, the outer membrane can be stiffer than the cell wall, and the mechanical pressures are often balanced between these structures {Rojas, 2018 #63}.

Numerous significant discoveries have recently emerged in the field of envelope biology. However, despite these advancements, there remains a substantial amount to unravel about this complex structure. We have yet to comprehend fully the assembly, regulation, and precise coordination of various functions within this multi-layered architecture.

We have a more extensive understanding of essential envelope factors compared to non-essential yet crucial elements in envelope biology. Our objective was to identify all non-essential genes necessary for maintaining for envelope integrity. Using the Gram-negative bacterial model organism *E. coli*, we developed a sensitive assay tailored to detect envelope integrity defects. In a high-throughput screening, we investigated a system ordered single-gene deletion library of *E. coli,* which encompassed 102 genes of unknown function in envelope biology. One of the most notable outcomes in our assay at low temperatures led us to focus on the gene *elyC* (<u>e</u>levated frequency of <u>ly</u>sis factor <u>C</u>, previously known as *ycbC*). Δ*elyC* mutant cells could be cultivated and maintained at 37°C without any apparent physiological defect. When sub-cultured at room temperature (21°C), Δ*elyC* mutant cells initially grew comparably to WT cells, but in late exponential phase, they exhibited a terminal phenotype of cell lysis in liquid media. We demonstrated that Δ*elyC* mutant cells had a block in PG assembly at 21°C, implicating *elyC*’s role in the proper metabolism and transport of the essential lipid carrier, undecaprenyl-phosphate, for the inner membrane step of the PG biosynthesis pathway {Paradis-Bleau, 2014 #10}. Subsequently, we discovered that the defective phenotypes of Δ*elyC* mutant cells could be alleviated by the overexpression of specific periplasmic chaperones. Furthermore, we found that Δ*elyC* mutant cells contained twice as many envelope protein aggregates as WT cells at 21°C, suggesting issues with envelope protein integrity and folding that result in a block in PG biogenesis when ElyC is absent at this temperature {Kouidmi, 2018 #67}.

The *elyC* open reading frame is predicted to encode a protein with two transmembrane helixes spanning the inner membrane, a small cytoplasmic loop and a large, highly conserved domain of unknown function as DUF218, located in the periplasm. Given the abundance and widespread distribution of DUF218 domains in bacteria, it suggests their vital role(s) in bacterial biology {Finn, 2008 #78; Paradis-Bleau, 2014 #10}.

In bacteria, particularly the Gram-negative *E. coli* K12 strain, approximately 35% of the synthesized proteins in the cytoplasm translocate and fold within the cell envelope or beyond it {Orfanoudaki, 2014 #90; De Geyter, 2016 #79}. The process of Protein trafficking and folding into their correct structures and locations are tightly regulated by complex and sophisticated mechanisms. An array of soluble, diffusible chaperones and folding catalysts facilitates the translocation of polypeptides and maintains their integration in the periplasm. The accuracy of these processes must be maintained under varying environmental conditions, including stress and temperature fluctuations. In cases of failure, proteases are mobilized to degrade mis localized or aggregated proteins {Bury-Moné, 2009 #78}{De Geyter, 2016 #79}. In *E. coli*, five primary pathways oversee protein folding under stress conditions {De Geyter, 2016 #79}. Consequently, a complex network and envelope stress response pathways assume responsibility for cell envelope biogenesis, maintenance, and antibiotic resistance.

We currently understand that the *E. coli* DUF218-containing protein ElyC is required for maintaining envelope protein integrity/folding and PG biosynthesis at a lower temperature of 21°C {Kouidmi, 2018 #67; Paradis-Bleau, 2014 #10}. However, the extent of ElyC’s role in envelope biology at the physiologically relevant temperature of 37°C remains unexplored. To gain deeper insights into our understanding of ElyC in envelope biology, we performed the phenotypic characterization and transcriptomic studies on the ElyC depleted strain grown at both 21°C and 37°C, comparing the results to those of WT cells.

We have demonstrated a significant reduction in PG content in the deletion mutant, with a 55% decrease at 21°C and 38% at 37°C. These findings align with our observations that the mutant cells undergo lysis at 21°C but not at 37°C. Furthermore, at 37°C, the Δ*elyC* cells exhibited increased sensitivity to β-lactam antibiotics, further confirming a PG-related defect. Additionally, our protein aggregation assay revealed a protein-folding defect in the *ΔelyC* mutant at 37°C, consistent with our prior findings{Kouidmi, 2018 #67}. In our RNA-Seq transcriptomic analysis, we identified a subset of genes that were activated in response to envelope stress at 21°C, with a more moderated response at 37°C. Specifically, 94 genes were upregulated at 21°C and 32 at 37°C, while 53 genes at 21°C and 19 genes at 37°C were downregulated upon ElyC depletion. The transcriptomic data highlighted the activation of stress response pathways such as Rcs, σ^E^, Cpx, and Bae, which are closely linked to PG biosynthesis and cell envelope biology, particularly involving oxidative stress, chaperone, or protease activities. Overall, this study underscores the broad physiological impact of ElyC to the bacterial cell envelope and PG biosynthesis pathway at optimal and suboptimal temperatures. These results validate our earlier findings regarding the extent of PG damage at different temperatures and confirm that the primary defect in the Δ*elyC* mutant lies within the envelope compartment. Additionally, our study sheds light on the consequences of eliminating ElyC function on the transcriptome and its impact on cell physiology.

## RESULTS

### ElyC-defective cells display increased width but maintain growth at 37°C

Our earlier research revealed that Δ*elyC* mutant cells exhibited pronounced phenotype defects in Luria-Bertani (LB) media at 21°C. They displayed slow growth and formed smaller colonies compared to WT cells on agar plates. In liquid media, Δ*elyC* mutant cells initially grew at a pace like WT cells, reaching the late-exponential phase (OD_600nm_ ≈ 0.6), but then ceased growing and underwent lysis {Paradis-Bleau, 2014 #10}. To investigate ElyC’s role in envelope biology at 37°C, we conducted a comparative analysis of the physiological characteristics of WT and Δ*elyC* mutant cells grown in solid and liquid LB media at both 37°C and 21°C.

We initiated a study by examining the growth and cell morphology of the Δ*elyC* cells at 21 vs 37°C. Cell growth was monitored in LB medium with 1% NaCl (see Materials and Methods). This salt concentration exacerbates the phenotypes of the Δ*elyC* mutant at 21°C, as previously demonstrated {Paradis-Bleau, 2014 #10. At, 21°C, as shown in Fig 1A, Δ*elyC* cells began to lyse around the late mid-log phase (OD_600_ ∼0.6). In contrast, at 37°C, the mutant cells exhibited growth as well as the wild-type strain and did not undergo lyse (Fig. 1B). Microscopic analysis of the cells at OD_600_ = 0.5 revealed no evident morphological defects in Δ*elyC* cells at 37°C (Fig. 1C), in contrast to pronounced bulging and lysis observed at 21°C. Furthermore, microscopic images taken at the late-exponential phase (OD_600_) showed that the Δ*elyC* cells continued to grow normally at 37°C (data not shown). Image analysis of the cells taken at OD_600_ = 0.5 disclosed a statistically significant (p < 0.01) thickening of the Δ*elyC* cells at 37°C compared to those of the wild-type grown at the same temperature (Fig. 1D). This particular phenotype of the Δ*elyC* cells grown at 37°C suggest a defect at the cell envelope level, particularly in PG biosynthesis. However, mutant cells exhibited similar growth to WT cells on solid media at 37°C. Microscopic analysis of the mutant cells revealed that the cells lysed occurred via membrane blebs. Notably, Δ*elyC* mutant cells displayed increased width compared to wild-type cells.

**FIG 1.**
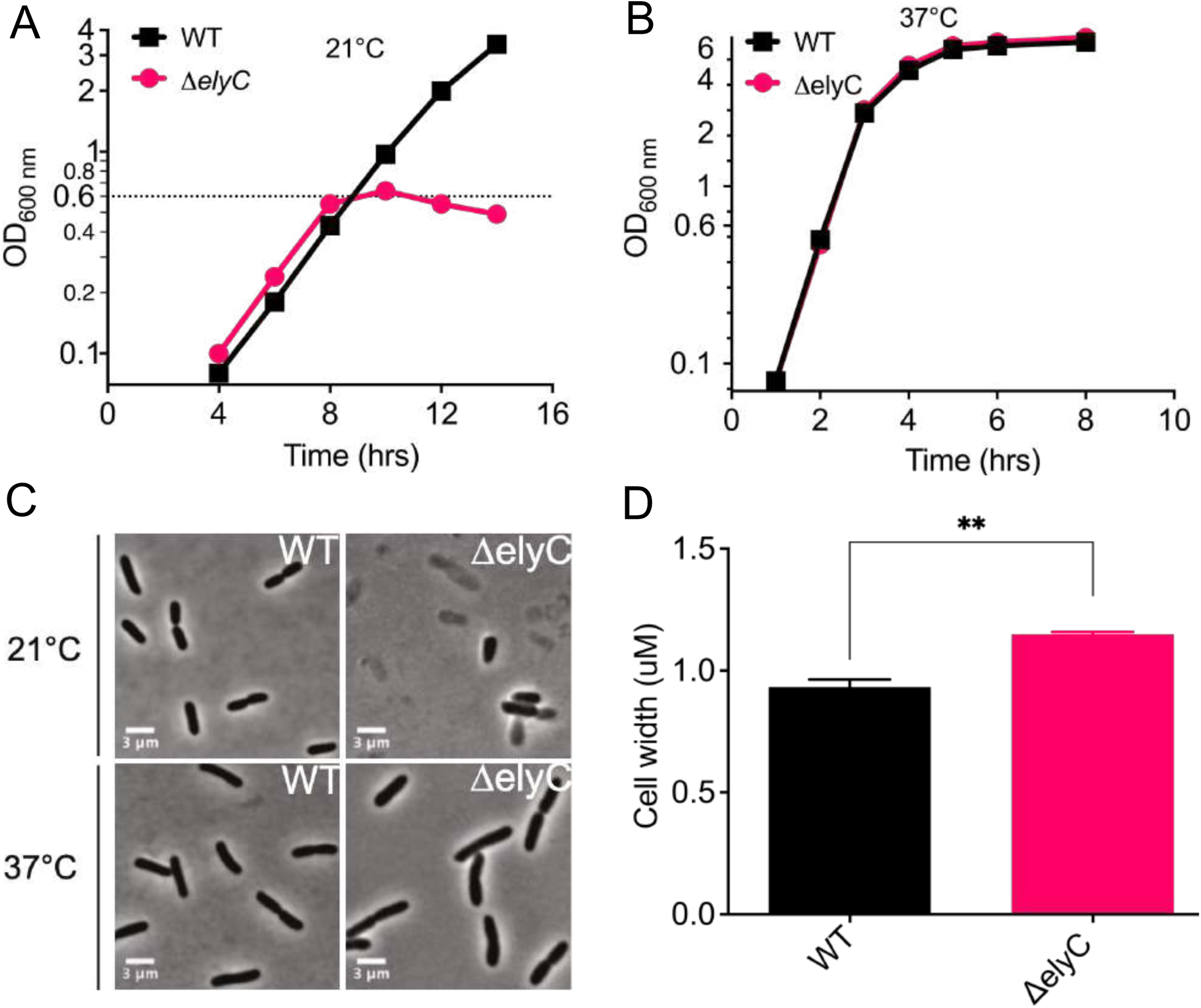
Deletion of ElyC induces cell lysis at 21°C and increased cell width at 37°C. Both *ΔelyC* mutant and wild-type cells were cultured overnight at 37°C, then diluted (OD_600_= 0.02) and grown in LB medium at 21°C and 37°C with continuous agitation (250 rpm). (A) *ΔelyC* mutant cells exhibit cell lysis at 21°C, indicated by growth inhibition around OD_600_ ∼0.6. (B) They display normal growth at 37°C. (C) Microscopic images at OD_600_= 0.5 show cell bulging and lysis in *ΔelyC* mutant cells at 21°C (upper right corner), with no morphological defects observed at 37°C (lower right corner). (D) *ΔelyC* cells exhibit a significant increase in cell width at 37°C compared to the wild type (p < 0.01, t-test). The width of 100 cells per replicate was measured and analyzed with Fiji software. Error bars represent mean ± SD; asterisks indicate significant differences (p-value < 0.05).

### PG biosynthesis defect in ElyC-depleted cells at 37°C

We previously demonstrated that PG biosynthesis is entirely obstructed in Δ*elyC* mutant cells after 2.5 generations at 21°C. Subsequently, mutant cells then continued to grow as well as WT cells for approximately 1.5 generations before ceasing growth and lysing, as documented {Paradis-Bleau, 2014 #10}. We proceeded to quantify the PG material precisely and found that Δ*elyC* mutant cells contain approximately 40% of the total relative PG amount of WT cells at OD_600nm_ of 0.5 {Kouidmi, 2018 #67}. The observed morphological difference in Δ*elyC* mutant cells grown at 37°C compared to WT cells suggests a substantial alternation in PG quantity and/or composition. This envelope layer is responsible for determining bacterial cell shape and maintaining this structure despite variations in osmotic pressure. To confirm the necessity of ElyC for proper PG biosynthesis at physiological temperature, we conducted a quantitative analysis of PG from both the WT and Δ*elyC* strains. PG sacculi were isolated from cells grown to an OD_600nm_ of 0.5 in liquid media at 37°C and 21°C, followed by digestion with muramidase. The resulting soluble muropeptide mixtures underwent analysis via ultraperformance liquid chromatography (UPLC) {Alvarez, 2016 #56}.

Relative PG quantities were calculated by comparing them to the total PG amount from WT cells grown at 37°C, serving as the reference sample. Our quantitative analysis reveals that Δ*elyC* mutant cells contain significantly less PG material than WT cells at both temperatures. The absence of ElyC leads to a reduction of about 40% in PG content at 37°C (Fig. 2A). This aligns with the observed cell shape defect in Δ*elyC* mutant cells grown at 37°C (Fig. 1C-D). Notably, *E. coli* cells can continue to grow normally with up to 50% less PG per cell {Prats, 1989 #7} and that Δ*elyC* mutant cells still maintain approximately 60% of the total PG material at 37°C (Fig. 2A), which explains their ability to grow similarly to WT cells at this temperature (Fig. 1B). The PG defect in Δ*elyC* mutant cells is more severe at 21°C than at 37°C. Mutant cells have only around 40% of PG material left once they reach an OD_600nm_ of 0.5 at 21°C, after which they cease growing and start lysing (Fig. 1A). Detailed muropeptide composition analysis did not indicate any significant difference in PG chemistry or structure between the samples (Fig. 2B-C, Table S1), implying a decrease in overall PG biosynthesis in Δ*elyC* mutant cells exacerbated at 21°C comparted to 37°C, rather than modification or degradation of PG.

**FIG 2.**
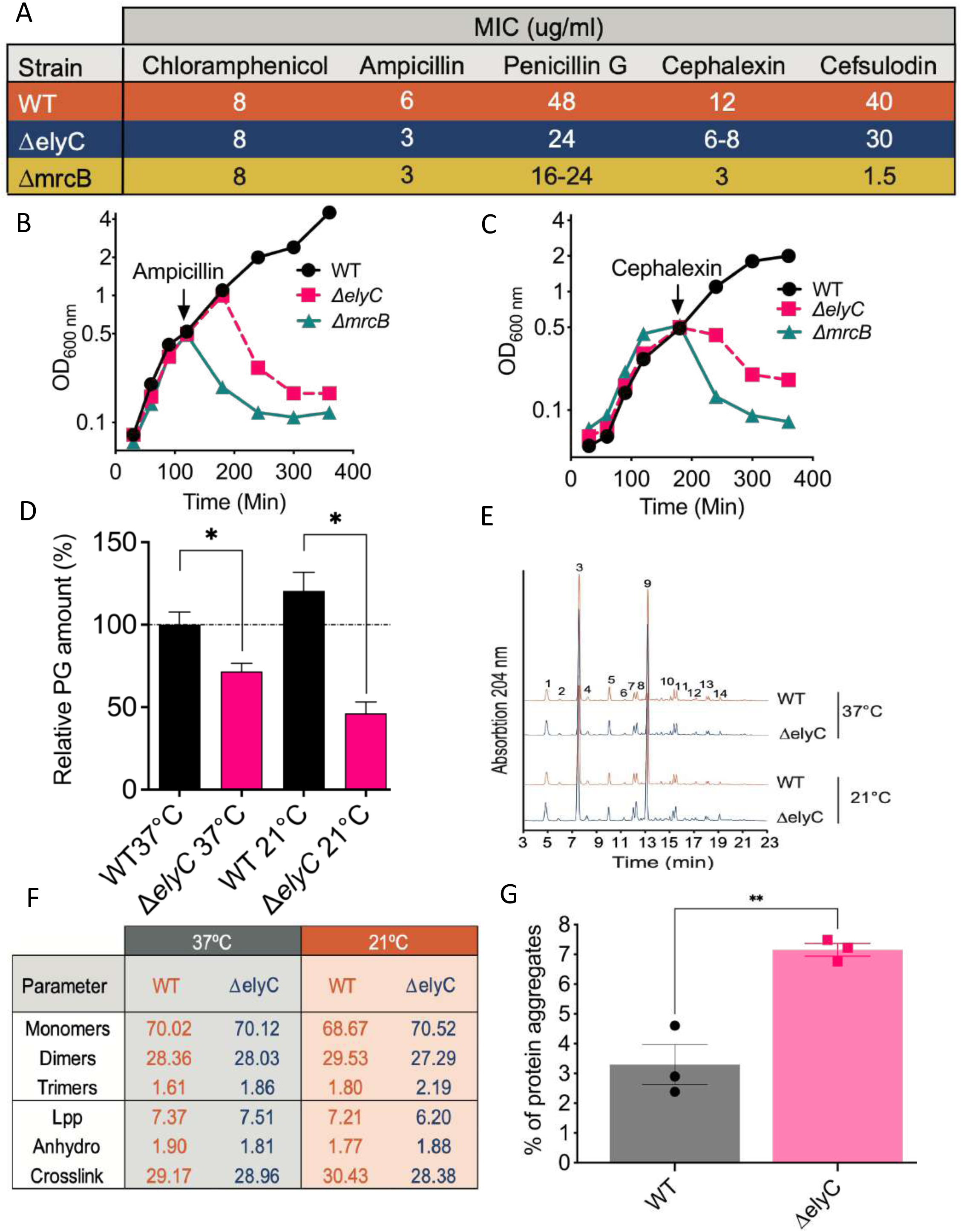
Sensitivity of *ΔelyC* mutant to *β*-lactam antibiotics at 37°C versus wild type, PG quantitative analysis and protein aggregation comparison in *E. coli* Cells. (A) Minimum inhibitory concentration (MIC) of *β*-lactam antibiotics for the indicated strains, determined from three independent experiments. (B, C) Lytic effect of *β*-lactam antibiotics. Cultures of WT, *ΔelyC*, and *ΔmrcB* were grown at 37°C to mid-log phase (OD_600_=0.5) and exposed to (B) Ampicillin (30 µg/ml) or (C) Cephalexin (30 µg/ml). The arrow indicates the time of treatment. (D) The PG amounts in *ΔelyC* mutant cells were significantly lower at both 37°C and 21°C compared to the wild type. PG purified from LB-cultured cells at OD_600_= 0.5, underwent UPLC analysis, with relative amounts calculated by comparing UPLC total intensity to the WT reference at 37°C. Statistical analysis involved three biological replicates, and error bars represent means ± SD, with asterisks indicating significance (p < 0.05). (E) Chromatogram displays muramidase-digested PG from WT and *ΔelyC* mutant cells at 37°C and 21°C. Identified muropeptide peaks are labelled with numbers, and details are provided in Table S1. (F) The graph displays relative molar abundance, including monomers, dimers, trimers, outer membrane lipoprotein-bound muropeptides (Lpp), and muropeptides with (1-6 anhydro) N-acetylmuramic acid (Anhydro). The percentages of cross-linkage are also provided. (G) Protein aggregation levels compared between *E. coli* wild type and *ΔelyC* mutant cells, determined from three experiments. Significance was assessed by unpaired t-test (Unequal Variances), with an asterisk indicating a significant difference (p < 0.005).

### ElyC deletion strain exhibits increased sensitivity to β-lactam antibiotics compared to WT cells at 37°C

If the Δ*elyC* mutant is indeed implicated in reducing PG levels at 37°C, it is reasonable to suspect that these mutant cells may exhibit increased susceptibility to β-lactam antibiotics, which specifically target the peptidoglycan biosynthesis process. To investigate this, we assessed the sensitivity of the Δ*elyC* mutant at 37°C to a range of β-lactam antibiotics, including ampicillin, penicillin G, cephalexin and cefsulodin, by measuring the minimum inhibitory concentrations (MIC). To provide a basis for comparison, we used a known β-lactam-sensitive strain, the penicillin-binding protein 1b (PBP1b) defective mutant (Δ*mrcB*) {Gutmann, 1986 #19}. As shown in Fig. 3A, the Δ*elyC* mutant exhibited twice the sensitivity to ampicillin and penicillin G compared to the wild-type strain, and appeared to be equaly sensitive to these antibiotics as the Δ*mrcB* mutant. In the case of cephalexin, the Δ*mrcB* mutant was even more sensitive (2.5-fold) than the Δ*elyC* mutant, which was itself almost twice as sensitive as the wild-type strain. The Δ*elyC* mutant displayed slight increased sensitivity to cefsulodin in comparison to the wild-type strain but notably more resistant than the Δ*mrcB* mutant. Confirming our expectations, the MIC for chloramphenicol, a protein synthesis inhibitor used as a control, remained consistent across all three strains. The sensitivity of both Δ*mrcB* and Δ*elyC* mutants to ampicillin and cephalexin was further evidenced by the observation of cell lysis when these antibiotics were added to log-phase (OD_600_ = 0.5) cell cultures (Fig. 3B and C). This cumulative data strongly supports the notion that the Δ*elyC* mutant exhibits a defect in PG synthesis at 37°C.

**FIG 3.**
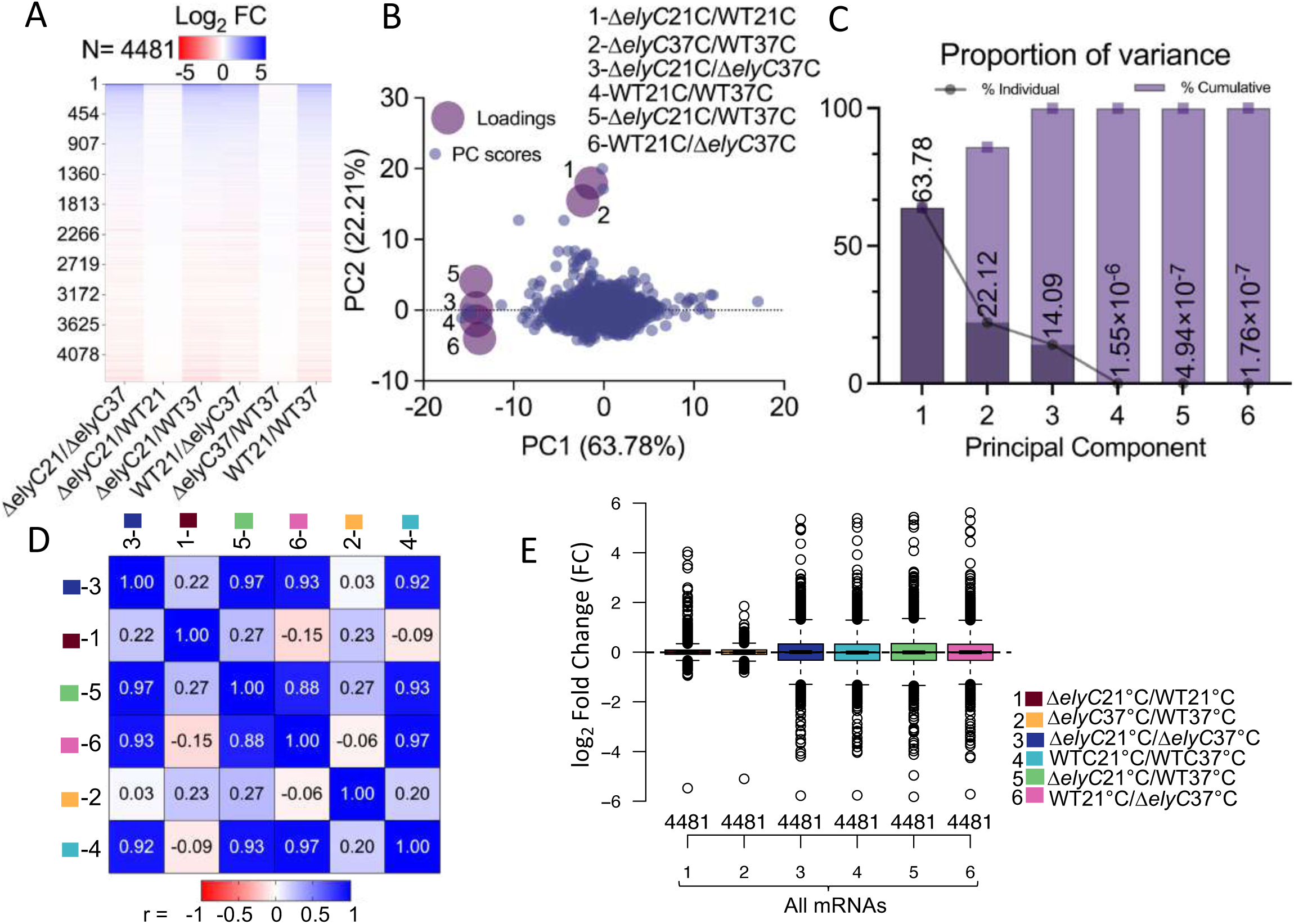
Diverse visualization methods show the impact of *ΔelyC* mutant versus wild-type strains and temperature shifts on gene expression. Heatmaps revealed distinct expression changes and grouping in conditions 1-*ΔelyC*21°C/WT21°C and 2-*ΔelyC*37°C/WT37°C compared to other comparisons. (B-C) PCA analysis indicated closer associations in comparisons 1, 2, and 3, contributing 63.78%, 22.12%, and 14.09% of the variance, while the other three conditions were closely grouped. (D) Pearson correlation showed high consistency in conditions 3, 4, 5, and 6, with lower correlation in comparisons 1 and 2, 1 and 3, and 2 and 3. (E) Notched box plots showed differences in log_2_ fold change values for the strain/mutant/condition across all mRNAs (cols. 1-6). Condition 2 had lower gene expression than condition 1, while conditions 3-6 exhibited similar expression patterns.

### Deletion of ElyC induces envelope protein integrity/folding defect at 37°C

Membrane-anchored lipoproteins and soluble periplasmic proteins constitute two distinct classes of proteins that must correctly fold within the periplasmic space to function effectively {De Geyter, 2016 #81}. Molecular chaperones play pivotal role in guiding proteins to attain their proper three-dimensional structure while preventing the aggregation of misfolded or damaged proteins {De Geyter, 2016 #81}{Tomoyasu, 2001 #78}. In a prior experiment, we demonstrated that overexpressing the periplasmic chaperones DsbG or Spy was sufficient to rectify the protein-folding defect and restore PG biosynthesis in ΔelyC mutant cells cultivated at 21°C. As a result, we conclude that the PG defect in the absence of ElyC is, at least in part, attributed to protein-misfolding {Kouidmi, 2018 #67}. Thus, we resonate to explore the possibility of protein aggregation in Δ*elyC* mutant cells grown at 37°C. To this end, we followed the protein aggregate isolation protocol outlined by Tomoyasu *et al*. in 2001 {Tomoyasu, 2001 #78}, with some modifications described in Materials and Methods section. This procedure enables the isolation of total cellular protein aggregates. Subsequently, we quantified both total and aggregated protein levels and conducted a t-test statistical analysis on data derived from three biological replicates. The results revealed that the level of aggregated proteins in the Δ*elyC* mutant was twofold higher than in the wild-type strain, constituting 7.15% and 3.29% of the total protein content, respectively (see Fig. 4). Therefore, our findings indicate an elevated occurrence of protein aggregation at 37°C in Δ*elyC* cells as compared to the wild-type strain.

**FIG 4.**
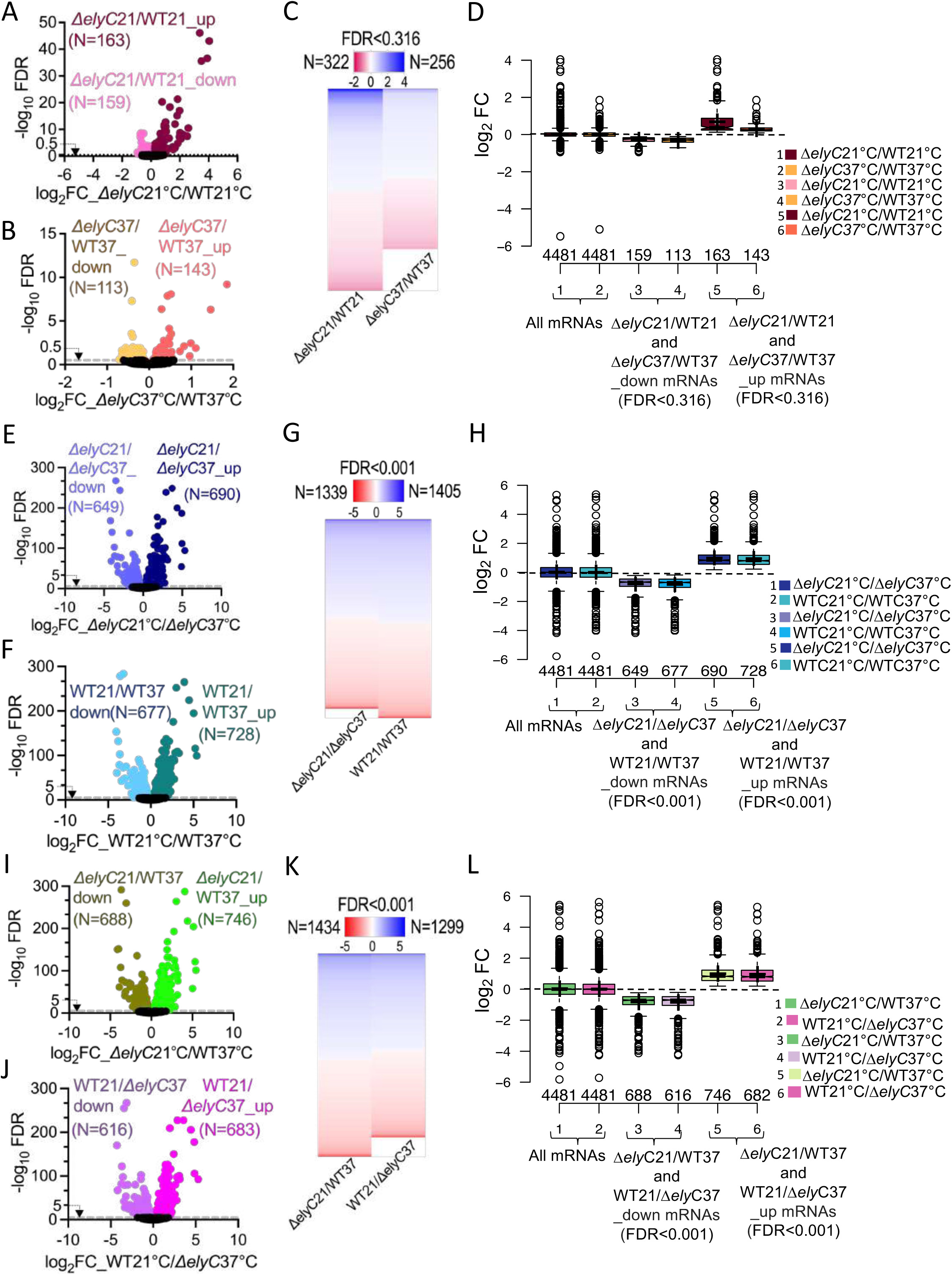
Elimination of elyC shows mRNA transcriptional changes between *ΔelyC* and WT cells at 21°C and 37°C. (A) Volcano plot depicting log_2_ ratios of 4480 mRNAs in *ΔelyC* mutant versus WT *E. coli* cells (Δ*elyC*21°C/WT21°C) at 21°C. The x-axis represents each log_2_ fold change mRNA and the y-axis shows –log_10_ of the FDR determined by DESeq2 analysis. Genes significantly upregulated (N=163_up) and downregulated (N=159_down) in *ΔelyC* mutant versus WT cells at FDR < 0.316 are plotted in maroon/magenta circles, while others are in black below the 0.5 of –log10 FDR threshold (4158 mRNAs). (B) Volcano plot similar to (A), comparing gene expression in *ΔelyC*37°C/WT37°C mutant and WT cells at 37°C for the 4480 mRNAs. Genes with a significant increase (143_up) or decrease (113_down) in WT cells at FDR < 0.316 are plotted in orange/yellow circles. (C) Heatmap analysis of the two comparisons for the 322 and 256 mRNAs (descended and arrayed) with significant gene expression log_2_ fold change values. The color scale ranges from 4 (dark blue) to –2 (dark red), showing overall higher gene expression patterns in *ΔelyC*21°C/WT21°C compared to condition ΔelyC37°C/WT37°C. (D) Notched box plots of total mRNAs in conditions 1 and 2 (D-cols 1 and 2) confirm the higher expression patterns in condition 1. Dissection of up and downregulated genes reveals a pronounced amount of upregulated genes in condition 1 (D-col. 5) versus condition 2 (D-col. 6), while downregulation shows no significant mean difference. (E) Volcano plot depicting log2 ratios of 4480 mRNAs in elyC mutant versus WT *E. coli* cells for *ΔelyC*21°C/*ΔelyC*37°C. Genes significantly upregulated (N=690_up) and downregulated (N=649_down) in elyC mutant versus WT cells at FDR < 0.001 are plotted in dark blue/light blue circles, while others are in black below the 5 of –log10 FDR threshold (3141 mRNAs). (F) Volcano plot similar to (E), comparing gene expression in WTC21°C/WTC37°C cells for the 4480 mRNAs. Genes with a significant increase (728_up) or decrease (677_down) in WT cells at FDR < 0.001 are plotted in dark green/light navy circles. (G) Heatmap analysis of the two comparisons for the 1339 and 1405 mRNAs (descended and arrayed) with significant gene expression log2 fold change values. The color scale ranges from 5 (dark blue) to –5 (dark red), showing overall higher gene expression patterns in WTC21°C/WTC37°C compared to condition *ΔelyC*21°C/*ΔelyC*37°C. (H) Notched box plots of all mRNAs in conditions 1 and 2 (H-cols 1 and 2) represent pretty close expression patterns in both conditions. Dissection of up and downregulated genes reveals the same amount of upregulated genes in both conditions (H-cols. 4-6) and shows no significant mean difference. (I) Volcano plot depicting log2 ratios of 4480 mRNAs in elyC mutant versus WT *E. coli* cells for *ΔelyC*21°C/WT37°C. Genes significantly upregulated (N=746_up) and downregulated (N=688_down) in elyC mutant versus WT cells at FDR < 0.001 are plotted in green/brown circles, while others are in black below the 5 of –log10 FDR threshold (3046 mRNAs). (J) Volcano plot similar to (I), comparing gene expression in WT21°C/*ΔelyC*37°C cells for the 4480 mRNAs. Genes with a significant increase (683_up) or decrease (616_down) in WT cells at FDR < 0.001 are plotted in dark pink/light pink circles. (K) Heatmap analysis of the two comparisons for the 1434 and 1299 mRNAs (descended and arrayed) with significant gene expression log2 fold change values. The color scale ranges from 5 (dark blue) to –5 (dark red), showing overall higher gene expression patterns in *ΔelyC*21°C/WT37°C compared to condition WT21°C/*ΔelyC*37°C. (L) Notched box plots of all mRNAs in conditions 1 and 2 (L-cols 1 and 2) represent close expression patterns in both conditions. Dissection of up and downregulated genes reveals the same number of upregulated genes in both conditions (L-cols. 4-6) and shows no significant mean difference.

### RNA-seq sample preparation, data processing, clustering, analysis, and visualization

To unravel transcriptomic variations in the ElyC-deleted strain compared to the WT strain at 21°C and 37°C, we conducted RNA sequencing. The cells were harvested during the early-log phase (OD_600_ = 0.3-0.35), as we observed that the *ΔelyC* mutant experienced cell lysis around OD_600_ = 0.5. RNA-seq libraries were prepared for three biological replicates of both strains at both temperatures. The cDNAs were sequenced using the Illumina HiSeq 2000 platform, resulting in a coverage range of 18 to 26 million reads per sample (Table S2). When total counts were normalized relative to library size, the RNA sequencing libraries displayed strong correlation coefficients (|r| > 0.94 to 0.98, p < 0.0001), indicating consistent preparation methods across biological replicates of *ΔelyC* mutants and WT cells at 21°C and 37°C (Fig. S1-S2). The hierarchical clustering of the samples further confirmed their separation based on temperature conditions and the two different strains (Fig. S3). We utilized the DESeq2 tool for the initial statistical analysis of RNA-seq data, including the computation of log2 fold change (log_2_-FC), p-values, and significant Benjamini-Hochberg adjusted p-values to identify differentially expressed genes. The MA plot generated using R package illustrates the distribution of log_2_-FC in overall gene expression, providing overall insights into differential expression patterns (Fig. S4).

To delve into the transcriptional changes influenced by strains and temperature conditions, we compared log_2_ expression levels between *ΔelyC* and WT strains at 21°C and 37°C, resulting in six comparisons (1-*ΔelyC*21°C/WT21°C, 2-*ΔelyC*37°C/WT37°C, 3-*ΔelyC*21°C/*ΔelyC*37°C, 4-WTC21°C/WTC37°C, 5-*ΔelyC*21°C/WT37°C, 6-WT21°C/*ΔelyC*37°C). The comparisons were visualized through heatmaps, Biplot (PCA), proportion of variance in PCA, Pearson correlation, and Box Plots to comprehensively explore the effects of strains and temperature shifts. The heatmap analysis on all mRNA shows that conditions 1-ΔelyC21°C/WT21°C and 2-ΔelyC37°C/WT37°C exhibited lower expression changes (Fig. 3-A) and appear to be closely grouped (Fig. 3-B), contrasting with the other four conditions. Upon closer PCA analysis, comparisons 1, 2, and 3 collectively contributed 63.78%, 22.12%, and 14.09% of the variance, indicating a closer association. Conversely, the other three conditions were closely grouped (Fig. 3-C). For additional insight, we used Pearson correlation to quantitatively assess linear relationships in gene expression profiles, enhancing our understanding of RNA-seq data patterns alongside heatmap and PCA. In our analysis, conditions 3, 4, 5, and 6 exhibited high consistency (r > 0.88), while comparisons 1 and 2, 1 and 3, and 2 and 3 showed lower correlation (r < 0.23).

Interestingly, conditions 1 and 4, 1 and 6, and 2 and 6 displayed negative correlations (r < –0.09), emphasizing distinct patterns in gene expression relationships (Fig 3-D). Notched box plots of log2 fold change were employed for all mRNAs to visually compare medians between groups and provide insights into the statistical significance of differences, as indicated by the notches representing 95% confidence intervals around the medians. Condition 2 exhibited lower gene expression compared to condition 1 (Fig 3-E, cols. 1-2) while conditions 3, 4, 5, and 6 displayed similar expression patterns with no significant differences in medians (Fig 3-E, cols. 3-6).

### Eliminating *E. coli elyC* revealed more mRNA changes in condition 1-*ΔelyC*21°C/WT21°C compared to condition 2-*ΔelyC*37°C/WT37°C

To assess the impact of *elyC* deletion on *E. coli* mRNA expression at 21°C and 37°C, considering the role of temperature and ElyC in gene regulation, particularly in peptidoglycan and outer membrane biogenesis, we filtered transcripts at FDR < 0.316 (Additional file 3) and generated a volcano plot (Fig. 4-AB). The smallest significant (FDR= 0.044004732) transcript reduction for condition 1(1-*ΔelyC*21°C/WT21°C) is 0.52-fold change (gene *arnC*), and the largest significant for this condition was 16.4-fold change, FDR (8.07488e^-44^) (gene *bdm*).

The smallest significant (FDR= 0.26) transcript reduction for condition 2 (2-*ΔelyC*37°C/WT37°C) is 0.61-fold change (gene *ynaK*), and the largest significant for this condition was 3.61-fold change, (FDR= 6.31444e^-10^) (gene *ymgG*). The smallest significant (FDR= 4.2054e^-169^) transcript reduction for condition 3 (3-*ΔelyC*21°C/*ΔelyC*37°C) is 0.05-fold change (gene *fimA*), and the largest significant for this condition was 41.16-fold change, (FDR= 7.13199e^-95^) (gene *ymgC*). The smallest significant (FDR=1.4167e^-284^) transcript reduction for condition 4 (4-WTC21°C/WTC37°C) is 0.11-fold change (gene *ompT*), and the largest significant for this condition was 41.76-fold change, (FDR= 3.9178E^-100^) (gene *ymgC*). The smallest significant (FDR= 3.5371e^-292^) transcript reduction for condition 5 (5-*ΔelyC*21°C/WT37°C) is 0.08-fold change (gene *sslE*), and the largest significant for this condition was 16.84-fold change, (FDR= 3.2174e^-288^) (gene *sra*). The smallest significant (FDR= 2.9933e^-268^) transcript reduction for condition 6 (6-WT21°C/*ΔelyC*37°C) is 0.11-fold change (gene *ompT*), and the largest significant for this condition was 11.89-fold change, (FDR= 3.4635e^-228^) (gene *sra*).

### ElyC deletion amplifies activation of numerous genes at 21**°C**, linked to envelope stress response and uncharacterized functions

Within the set of upregulated transcripts at 21°C, 61 genes were identified based on an expression ratio of ≥ +1.4 FC cut-off threshold, and these were grouped into five distinct functional categories. These functional categories encompass envelope-induced response genes (44.27%), genes with unknown functions (37.7%), regulatory elements (13.11%), oxidative stress-related genes (3.28%), and genes involved in protein translation (1.64%). The percentage of genes expressed within each functional category provides a comprehensive overview of the genome’s response in the Δ*elyC* mutant under sub-optimal temperatures. A closer examination of the functional category of envelope-induced response genes at 21°C in the *ΔelyC* mutant reveals that the majority of these genes are associated with process such as extracellular polysaccharide formation, colanic acid biosynthesis, transferase activity, osmotic stress response, PG protection, cardiolipin phospholipid biogenesis, and periplasmic protein folding.

Another prominently represented group of upregulated genes belongs to the category of unknown function. Notably, the transcriptional upregulation of genes like *ydeI, yggE,* and *yghA,* which encode predicted envelope proteins involved in oxidative stress defence and oxidoreductase activity was observed. It is worth noting that the expression fold change of *yggE* exhibited a lower value (1.27). The *yggE* transcription is known to be upregulated by the general σ^S^ stress response factor {Peano, 2015 #26}, and a previous study indicated that *ydeI* is upregulated following the envelope stress response {Lorenz, 2016 #31}. The regulated genes related to peptidoglycan were not highly pronounced; however, the Δ*elyC* mutant under sub-optimal temperature exhibited a significant, albeit partial reduction in the expression of the peptidoglycan biosynthesis genes such as *murE* {Michaud, 1990 #32}, as well as hydrolysis genes *mepM* and *pbpG* {Romeis, 1994 #33; Singh, 2012 #34} (Table S3).

Our transcriptomic analysis uncovers that the upregulated genes in the Δ*elyC* mutant at 21°C are predominantly associated with the presence of a highly defective cellular envelope. This is evident from the activation of several envelope stress response systems and their related genes. This response underscores that the susceptibility of the Δ*elyC* mutant {Paradis-Bleau, 2014 #47}at 21°C is primarily linked to defects within the envelope compartments, potentially resulting from peptidoglycan damage, among other factors.

### ElyC deletion leads to downregulation of multiple genes at 21°C, primarily impacting cell envelop, motility, regulatory, and membrane transporter functions

In the set of downregulated transcripts at 21°C, 22 genes were identified based on an expression ratio of ≤ –1.4 FC cut-off threshold, and they were classified into different functional categories. These categories include cell motility (45.45%), membrane transporters (18.19%), genes with unknown functions (13.63%), regulatory elements (13.63%), and outer membrane biogenesis (9.1%) (Fig. 6A, and Table 2). The percentage of genes expressed within each functional category offers a comprehensive assessment of the genome’s response in the Δ*elyC* mutant under sub-optimal temperature. Based on the defined cut-off ratio, we observed that the most downregulated genes are primarily associated with flagellar motor activity, membrane transporters, functions of unknown nature, regulatory elements, and outer membrane biogenesis. However, before applying the cut-off threshold, the most notably downregulated genes in the Δ*elyC* mutant at 21°C are primarily involved in specific cellular envelope functions (membrane transporters), motility, and outer membrane biogenesis (Table S4). Several of the downregulated genes play roles in high-energy-consuming processes, such as cell mobility and protein-export machinery. For instance, various flagella biosynthesis genes, including the key regulators of flagellum genes expression *flhDC*, as well as essential components of the protein export machinery system, like the ATPase SecA and accessory proteins SecD, SecF, and YidC, display partial downregulation (Table 2 and Table S4). Additionally, we observed the downregulation of genes related to cellular transport systems (Table 2), encompassing amino acid and sugar transport, mechanosensing channels, and multidrug efflux systems. Many of these transporters rely on energy derived from the proton motive force or ATP hydrolysis, mainly produced during the respiratory process {Haddock, 1977 #38; Jormakka, 2003 #39}. Furthermore, genes like *mlaF, mlaE, mlaD, mlaC,* in the Δ*elyC* mutant at 21°C, which are involved in outer membrane biogenesis, also exhibit significant, albeit partial reduction in gene expression when filtered by the ≤ –1.4 FC cut-off threshold (Table S4). These genes from components of the ABC transporter are responsible for maintaining outer membrane asymmetry. Notably, important components of the aerobic respiratory chain, including the cytochrome C-encoding gene *cyoC,* subunit I of the cytochrome *bo* terminal oxidase complex (*cyoB*), and the electron-feeding respiratory chain Gcd, are also partially downregulated, (Table S4). Genes associated with lipid A modification, such as *arnC* and *eptA*, along with their signal activator in the two-component system BasSR {Shimada, 2012 #40}, are observed to be downregulated (Table 2). The functional categorization of the downregulated genes in the Δ*elyC* mutant at 21°C highlights their involvement in critical cellular processes, particularly in cell motility and envelope compartments, primarily within the inner membrane (IM) processes that either produce (respiratory chain) or require (transporters, motility or protein export) energy.

### Deletion of ElyC triggers fewer activations of envelop-induced response genes at 37°C

In the absence of ElyC at 37°C, the number of upregulated gene subsets was considerably reduced compared to 21°C. (Table S5). Specifically, we identified 14 genes that met the expression ratio threshold of ≥ +1.4 FC, which were classified into four functional categories. These categories encompass cell metabolism (35.71%), genes with unknown functions (35.71%), membrane transporters (14.9%), and envelope stress response (14.9%) (Fig. 6B, and Table 3). Transcription profiles of the Δ*elyC* mutant grown at 37°C reveal the upregulation of genes such as *wbbH* (1.45 FC), *degP* (1.4 FC), *hdhA* (1.26 FC), *mscL* (1.19 FC) and *bepA* (1.17 FC), all of which are involved in envelope stress response systems (Table 3, and Table S5). DegP and BepA function as periplasmic proteases and chaperones. MscL acts as a high-conductance mechanosensitive channel protein, HdhA is an osmotic stress-induced protein, and WbbH serves as an O-antigen polymerases. Within the group of genes with unknown functions and envelope-induced responses, *degP*, *ymgD*, and *ymgG* were also found to be upregulated both at 21°C and at 37°C (Fig. 6Ca). Additionally, we identified some upregulated genes related to ion efflux systems involved in cellular transport. In summary, the analysis of the upregulated genes at 37°C suggests that, unlike cells grown at 21°C, the envelope stress response is only minimally activated at 37°C.

### ElyC deletion also downregulates cell envelope biogenesis genes at 37°C

Functional groups of downregulated genes in the Δ*elyC* mutant at 37°C closely resemble those identified at 21°C (Fig. 6B, and Table 4). Notably, these group include genes related to cell mobility, particularly flagella apparatus genes. Genes such as the flagellum-specific ATP synthase FliI and the flagellar motor switch protein FliG exhibit downregulation at both 21°C and 37°C (Fig. 6Cb). Within the group of membrane transporters, the downregulated genes are primarily involved in ion exchange between the cytoplasmic and periplasmic compartments. Examples include the potassium translocating ATPase subunit KdpB, the sodium/proton antiporter NhaB, and the high-permeability porin OmpF. Furthermore, genes associated with components of the respiratory process are downregulated at both temperature conditions. At 37°C, the NADH dehydrogenase subunits NuoM/N experience reduced transcriptional levels. These subunits are vital for proton translocation during the respiratory process, leading to proton motive force (PMF) production {Friedrich, 1998 #41; Steimle, 2011 #42}. Additionally, in the group of protein export machinery, we identified the downregulation of the membrane protein insertion efficiency factor YidD and the lipoprotein maturation factor Lnt. The outer membrane biogenesis gene *mlaF* is downregulated both at 21°C and 37°C (Table S6). The *mlaF* gene codes for the ATP-binding subunit of the Mla phospholipid ABC transporter complex, which is involved in maintaining outer membrane asymmetry. The analysis of downregulated genes in the Δ*elyC* mutant suggests that cellular functions are impacted at 21°C and 37°C.

### Transcriptomic analysis reveals quality control pathways modulating protein folding in ElyC-defitient mutant at 21°C and 37°C

In *E. coli*, at least five pathways, namely Bae, Cpx, Psp, Rcs, and the sigma factors *rpoS* (σ^S^) and *rpoE*, are triggered in response to envelope stress {De Geyter, 2016 #79}{Bury-Moné, 2009 #78}. Under certain conditions, multiple pathways can become activated, and some genes exhibit functional redundancies while operating within interconnected networks. Our transcriptomics data analysis revealed that in an ElyC-depleted strain at 21°C, at least four stress response systems – σ^E^, Rcs Cpx, and Bae – were activated, whereas at 37°C, only Rcs and Cpx pathways were triggered. Details on the subset of genes and functional categories involved in these pathways are provided in Table 5.

Among the genes upregulated at 21°C, we identified specific genes, such as *otsA*, *otsB,* and *osmE*, that are associated with the activation of the general stress response factor σ^S^ {Giaever, 1988 #25; Peano, 2015 #26}. The downregulation of ElyC led to the decreased expression of *proW, proX, and proV* genes at 21°C, which are significantly influenced by σ^E^, and play a role in membrane transporter activity. Additionally, the *yafK* gene, whose expression is significantly modulated by the Cpx pathway, experienced downregulation and has an unknown function. Our study also revealed the upregulation of genes related to the Rcs pathway, including the transcriptional regulator *rcsA*, the osmotic stress protein OsmB, and genes associated with colanic acid biosynthesis (*wza, wzc, wzb, wcaA, wcaG*) {{Bury-Moné, 2009 #78}Laubacher, 2008 #20;Navasa, 2013 #21}, along with the periplasmic protein Ivy, which protects the peptidoglycan from lysozyme’s degradation {Deckers, 2008 #22}. Furthermore, genes related to the activation of Cpx pathway, namely *yceJ, yceI, degP, spy, ppiA, and hdhA* were upregulated (Table 5) {Vogt, 2012 #91}{Raivio, 2013 #24; Raivio, 2014 #23; Kouidmi, 2018 #67}. The absence of ElyC enhanced the activation of several genes at 21°C, whose functions are still unknown and whose expression is significantly modulated by RcsB and Cpx pathways.

The periplasm of *E. coli* harbor stress-response mechanisms, including chaperones like SurA, and DegP {Humes, 2019 #89}{De Geyter, 2016 #79}. Molecular chaperones act on non-native proteins in the cell, preventing their aggregation, premature folding, or misfolding, with distinct effects {De Geyter, 2016 #79}{Huang, 2016 #87}. After the synthesis of proteins within the cytoplasm, the secretory proteins must translocate to the plasma membrane via the Sec translocase system {Sardis, 2017 #88}. They should properly fold only at their final destinations or be degraded if mislocalized or aggregated by proteases activity {De Geyter, 2016 #79}. The unique features of secretory proteins have a significant impact on our understanding of protein trafficking, folding, and aggregation {Tsirigotaki, 2018 #86}. Additionally, some chaperones and proteases modulate protein folding/aggregation within the cell envelope under stress conditions. Our transcriptome profiling revealed the upregulation of genes encoding chaperones and protease activities. Specifically, we found upregulation of the *ivy, spy, degP*, *ppiA*, which play crucial roles in facilitating the transition of unfolded proteins during cytoplasmic transit and correct folding within periplasmic resident proteins at both 21°C and 37°C (Tables 1, 3, and 5). These chaperone genes are modulated by the Rcs, Cpx, and Bae pathways. Notably, *degP* was overexpressed at both 21°C and 37°C, highlighting the significant role of the DegP chaperone protein in ElyC-depleted cells at optimal and suboptimal temperatures. DegP is known to be specifically induced by the Cpx envelope stress response system {Raivio, 2013 #24;Raivio, 2014 #23} and functions as both a chaperone and protease for exported protein substances in the periplasm and outer membrane proteins {De Geyter, 2016 #79}. Furthermore, our study identified the transcriptional activation of a small RNA *omrA*, *omrB* {Brosse, 2016 #27; Guillier, 2006 #28}, and the periplasmic protein folding catalyzer *fkpA* {Humes, 2019 #89}{Dartigalongue, 2001 #29; Danese, 1997 #30}, which are associated with the EnvZ/OmpR two-component system and the envelope stress response factor σ^E^, respectively.

## DISCUSSION

Significant progress has been made in understanding peptidoglycan (PG) biosynthesis mechanisms, although certain steps, particularly those related to the inner membrane, remain poorly defined ElyC is a member of the widely conserved protein family with the DUF218 domain. Despite the prevalence of DUF218 domain-containing proteins in various bacteria, their precise functions, including that of ElyC, remain elusive {Finn, 2008 #78; Paradis-Bleau, 2014 #10}. While our prior research has demonstrated ElyC’s importance in PG biosynthesis at low temperatures {Kouidmi, 2018 #67}{Paradis-Bleau, 2014 #47},it remains unclear whether ElyC plays a role only at suboptimal temperatures or if it has distinct functions at both suboptimal and optimal temperatures. In this study, we aimed to elucidate the role of ElyC in PG biogenesis in Δ*elyC* and wild-type strains grown at 21°C and 37°C. We employed a combination of the phenotypic assessment (PG analysis and sensitivity to β-lactam antibiotics) and transcriptomics approaches to unravel ElyC’s significance in PG biosynthesis and the global gene expression pattern under ElyC depletion conditions in *E. coli* compared to the wild-type strain. For this experiment, cells were grown and harvested in triplicates, following established procedures. Our findings indicate that the Δ*elyC* mutant exhibited a defect in PG content and sensitivity to β-lactam antibiotics when grown at 37°C with more pronounced PG synthesis impairment observed at 21°C (Figs. 1-3), At an OD_600_ of ∼0.6, the Δ*elyC* mutant lysed at 21°C, whereas it continued to grow exponentially at both temperatures (Fig. 1-A-B-C). Notably, the absence of ElyC did not induce apparent morphological alternations in Δ*elyC* mutant cells grown at 37°C (Fig. 1-C). However, there was an increase in cell width (Fig. 1-D), suggesting that these cells had reduced resistance to internal pressure, resulting in increased cell width. After ∼ 5.5 hours of growth at low temperature, the Δ*elyC* mutant exhibited a PG assembly arrest and subsequently underwent lysis, one doubling time following the initial appearance of the PG defect. Our previous study, did not analyze PG assembly in Δ*elyC* mutant cells grown at 37°C {Paradis-Bleau, 2014 #10}. Our complementary peptidoglycan quantitative analysis revealed a clear defect in PG content at 37°C. The Δ*elyC* mutant strain had approximately 38% less PG at 37°C and less than 55% at 21°C compared to the wild-type strain. Remarkably, the structure and composition of the cell wall remained unaltered in the Δ*elyC* mutant, irrespective of the temperature tested. This suggests that a reduction in PG content, as long as it does not exceed 50%, does not affect cell morphology and growth {Prats, 1989 #43}. Therefore, it appears that the Δ*elyC* mutant at 37°C contains sufficient PG to maintain normal cell morphology and growth, whereas at 21°C the drastic reduction in PG content (55 %) leads to cell lysis. The substantial increase in cell width in the Δ*elyC* mutant at 37°C can be attributed to the decreased PG content at this temperature, implying that reduced PG content at 37°C may decrease cell rigidity and thickness, potentially widening the Δ*elyC* mutant cells due to the turgor pressure exerted on the inner membrane.

A study by Turner et al. (2018) {Turner, 2018 #68} demonstrated that altered cell shape is associated with significant changes in peptidoglycan biophysical properties. They combined atomic force microscopy with size exclusion chromatography and found that when *E. coli* cells adopt a spheroid shape, glycans become shorter and disordered. Such shape changes can result from external or internal factors like chemical or genetic inducements {Turner, 2018 #71}.

As the PG analysis did not show any differences in peptidoglycan composition or structure between the Δ*elyC* mutant and the wild type at both temperatures, likely, that the reduction in PG content is not a consequence of abnormal activation of PG hydrolytic or other modifying enzymes. Instead, it may result from a blockade in the PG biosynthesis pathway, particularly at the level of the inner membrane, possibly affecting the undecaprenyl phosphate lipid II carrier step {Paradis-Bleau, 2014 #10}. Furthermore, our data showed that the Δ*elyC* mutant was more sensitive to β-lactam antibiotics at 37°C compared to the wild-type strain. These antibiotics target the later stages of the peptidoglycan assembly pathway {Kohanski, 2010 #67}, suggesting that the cold-sensitive phenotype might be due to physiological changes occurring at low temperatures, such as modification in inner membrane phospholipid composition that could slow down or entirely block PG biosynthesis.

In our prior study, we demonstrated that the Δ*elyC* mutant’s defective protein folding led to increased levels of cell membrane protein aggregation and, ultimately, PG biosynthesis defect when cells were grown at 21°C. This defect could be rescued by overexpression of either DsbG or Spy genes {Kouidmi, 2018 #67}. To assess the possibility of protein aggregation in Δ*elyC* mutant cells grown at 37°C, we conducted a protein aggregation experiment at this temperature. We observed that the level of aggregated proteins was twofold higher in the Δ*elyC* mutant than in the wild type, resembling what was observed in the Δ*elyC* strain grown at 21°C. This suggests that interruption in PG biosynthesis in the Δ*elyC* mutant cells at 37°C may follow a similar mechanism of envelope protein folding defects as observed in the Δ*elyC* mutant cells grown at 21°C, albeit with distinct characteristics.

To gain deeper insights into the molecular mechanisms of PG biosynthesis and to obtain a comprehensive view of the gene expression in ElyC-depleted strains, we performed transcriptomic analysis and compared the results with the phenotypic assays. Our transcriptomic analysis revealed that ElyC deficiency at 21°C resulted in a greater number of differentially expressed genes compared to 37°C (Fig. 5A-B), indicating a relationship between the severity of the Δ*elyC* phenotypes and the magnitude of the transcriptional response.

**FIG 5.**
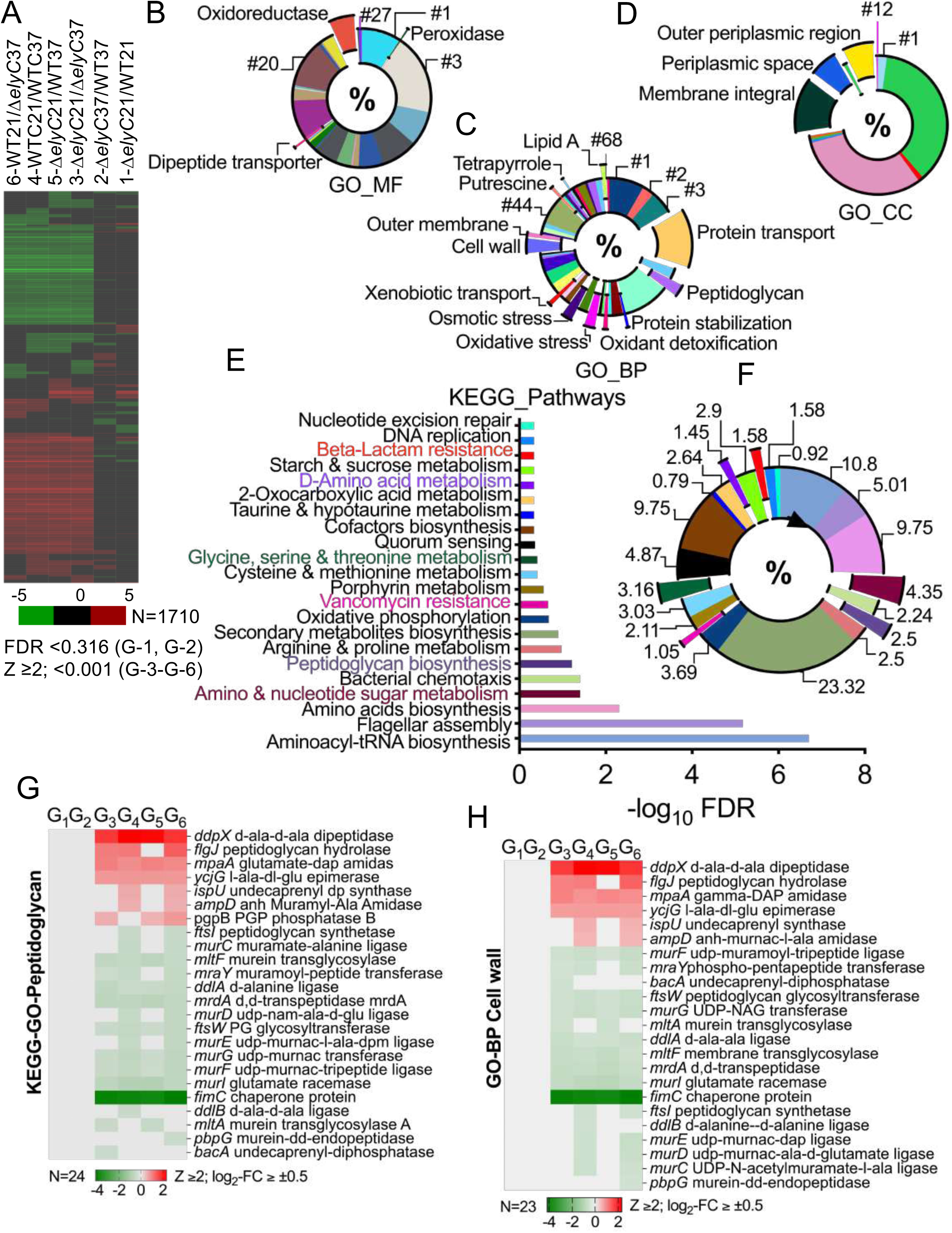
A comprehensive analysis identified 1710 differentially expressed mRNAs across six comparison groups (G1-G6). Utilizing heatmaps, GO enrichment analysis, and KEGG pathway analysis, distinct mRNA expression patterns were revealed in *ΔelyC* mutant cells grown at 21°C and 37°C compared to the WT strain, impacting specific biological processes, molecular functions, and cellular components. (A) Heatmaps of differentially expressed genes in *ΔelyC* mutant and wild-type (WT) cells reveal distinct mRNA transcriptional changes in G1 and G2 compared to the other four comparison groups. Initially, 1867 genes (supplementary File S3) were ranked using FDR cut-offs (<0.316 for G-1 and G-2, and <0.001 for G-3 to G-6) and visualized through a volcano plot (see Fig. 4). Subsequent analysis was performed using the BV-BRC transcriptomics tool with Z-score (Z ≥2), resulting in a refined mRNA set of 1710 entries (supplementary File S3) organized into heatmaps. The log2 fold change values are depicted on a color scale ranging from –5 (green) to 5 (red), where down-regulated genes are shown in green, up-regulated genes in red, and genes with no significant change are represented in black. (B-D) Donut charts depict the enrichment of GO terms related to molecular function (MF), biological process (BP), and cellular component (CC). Each donut chart highlights the most prominent feature relevant to the experimental context. Detailed information for each dataset is in supplementary File S4. (E-F) KEGG pathway analysis results illustrate different affected KEGG pathways in *ΔelyC* mutant cells compared to the WT strain across all six group comparisons. The most significant pathways are distinguished by different colors, and additional details about corresponding genes within each pathway are presented on the y-axis based on their –log10 FDR statistical analysis, as shown on the x-axis in Figure E. The percentage of each KEGG pathway is shown in Figure F. Detailed information for each dataset is in supplementary File S5. (G-H) Heatmaps highlight transcriptionally upregulated and downregulated genes involved in peptidoglycan biosynthesis, enriched in the KEGG pathway (FIG 5-E-F), and cell wall biosynthesis, enriched in GO-BP (FIG 5-C) defining representative functional categories across six groups (G1-G6) in the comparison. Values are displayed for log2-fold change (log2-FC) ≥ ±0.5 and Z-score (Z) ≥ 2, using a color scale ranging from –4 (green) to 2 (red), with down-regulated genes shown in green, up-regulated genes in red, and genes with no significant change represented in silver.

The RNA sequencing data demonstrated downregulation of certain genes at 21°C (Table 2), primarily at the inner membrane level, suggesting that the absence of ElyC may impact the cellular compartment and subsequently disrupt PG synthesis. If this interpretation holds, the flow-through of PG synthesis might be hindered or entirely blocked.

The downregulated genes in the Δ*elyC* mutant at 21°C included those involved in the electron transport chain, energy-consuming transport, flagella mobility, outer membrane biogenesis, and regulatory elements (Table 2). Notably, there was a slight downregulation of *cyoB* (−1.20 FC) and *cyoC* (−1.20 FC) genes (Table S4), which encode for subunits of the cytochrome *bo* terminal oxidase of the respiratory chain. This downregulation could be expected to reduce the respiratory process, potentially, affecting the ATP production and proton motive force (PMF). Notably, a similar transcriptional reduction of genes involved in the respiratory chain was observed at 37°C. Two other genes, *nuoM* (−1.35 FC) and *nuoN* (−1.31 FC), which code for subunits of NADH dehydrogenase, were also slightly downregulated at 37°C may be linked to the activation of the Cpx two-component system, as previous research shown that the Cpx response in *E. coli* can decrease the expression of genes associated with different components of the respiratory chain {Bury-Mone, 2009 #45;Raivio, 2013 #24}. Furthermore, genes encoding key component of protein export machinery, including SecA, SecF, SecD, and YidC, were slightly downregulated (Table S4) in the Δ*elyC* mutant at 21°C. Interestingly, it has been observed that mutations reducing the levels of Sec factors (Sec D, F) result in cold-sensitive mutants (cs), which are crucial for protein translocation and inner membrane protein insertion {Nouwen, 2005 #46}. *E. coli* mutants with individual *sec* gene deletions or mutations are unable to grow at low temperatures but exhibit normal growth at 37°C {Gardel, 1987 #47;Gardel, 1987 #47;Nouwen, 2005 #46;Pogliano, 1993 #48}. The altered composition of inner membrane phospholipids at low temperatures makes the protein export machinery less effective and more susceptible to mutations at this level. This leads to the accumulation of abnormal envelope proteins and the activation of the Cpx and RpoE extra cytoplasmic stress systems {Shimohata, 2007 #49}. In the Δ*elyC* mutant, the decrease in the transcriptional levels of *secA*, *secF*, *secD*, and *yidC* may contribute to the Δ*elyC* cold-sensitive phenotype and the activation of the Cpx and RpoE stress systems. Investigating this hypothesis in future studies would be of great interest.

Our RNA sequencing data also revealed upregulated genes at 21°C, with many of them having unknown functions and being associated with envelope-induced responses (Table 1). A comparison of these unknown function genes with those previously reported to be upregulated after treating cells with inhibitors of lipoprotein transport or antibiotics targeting peptidoglycan biosynthesis showed a significant overlap {Laubacher, 2008 #20; Lorenz, 2016 #31}, suggesting that many of these genes are linked to the envelope-induced response.

Additionally, we observed the activation of highly expressed proteins YmgG and YmgD (14.95 and 11.31 FC, respectively) with unknown functions (Table 1). These proteins have been reported to be induced in the presence of an envelope defect {Laubacher, 2008 #20; Lorenz, 2016 #31}. The level of activation of the envelope-induced response correlated with the severity of the peptidoglycan defect in the Δ*elyC* mutant.

Furthermore, we found that only three genes with chaperone/protease activities (DegP), and unknown function (YmgD, and YmgG) were upregulated, while two flagellar biosynthesis proteins were downregulated at both 21°C and 37°C (Fig. 6Ca). In contrast, more chaperone proteins such as Ivy, Spy, DegP, OmrA, OmrB, PpiA, and stress response pathways, including, Rcs, σ^E^, Cpx, and Bae {De Geyter, 2016 #79}{Bury-Mone, 2009 #28}{Vogt, 2012 #91}were activated in ElyC-depleted strain at 21°C compared to 37°C (Table 5). The overexpression of the periplasmic chaperone Spy, known for its effective ATP-independent chaperone activity in suppressing protein aggregation and aiding protein refolding {De Geyter, 2016 #79}{Quan, 2011 #89}, suggests a possible induction of the Bae system, as Spy is known to be induced by both the Cpx and Bae envelope stress systems {Srivastava, 2014 #44;Kouidmi, 2018 #100}. Therefore, we conclude that ElyC may have a temperature-dependence role in managing general or specialized chaperone factors proteins folding in the cell envelope of *E. coli*.

**FIG 6.**
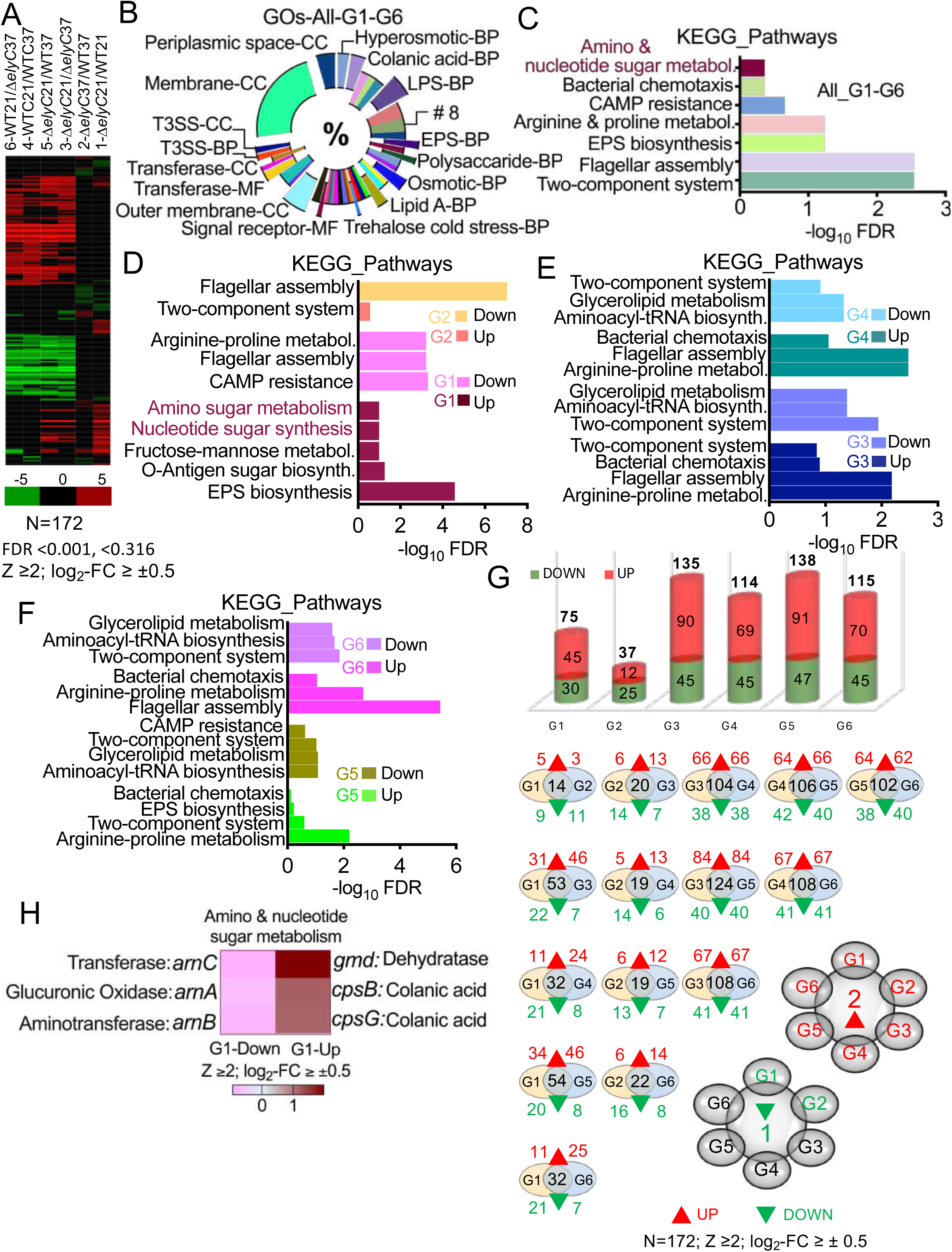
Analysis of 172 differentially expressed mRNAs (FDR <0.001, <0.316, Z ≥2; log2-FC ≥ ±0.5) in *ΔelyC* mutant vs WT across six comparison groups (G1-G6) using. (A) Heatmaps represent the expression patterns across the six comparison groups (G1-G6; Supplementary File S6). The log_2_ fold change values are color-coded from –5 (green) to 5 (red), with down-regulated genes in green, up-regulated genes in red, and unchanged genes in black. (B) Donut Plots illustrate the enrichment of GO terms (MF, BP, CC) for all groups (G1-G6), highlighting the most relevant features. Detailed data is in supplementary File S7. Individual up and down-regulated GO analysis is presented as Bubble Plots in supplementary Figs (FIG 6— Figs S5-S7), and detailed data is in supplementary Files 8-9. (C-F) KEGG pathway analysis results show affected pathways in *ΔelyC* mutant cells compared to the WT strain across all six comparison groups (C), and individually for up/down-regulated genes (D-F). The most significant pathways, such as amino sugar metabolism and nucleotide sugar synthesis related to peptidoglycan biosynthesis, are highlighted in red. Pathways are displayed based on their –log10 FDR statistical analysis (x-axis) in Figs. C-F. Detailed data is in supplementary Files S10 and S11. (G) Histograms illustrate the varying numbers of significantly expressed mRNAs across six comparison groups, with groups G1 and G2 showing lower counts compared to G3, G4, G5, and G6. Venn diagrams depict the differentially expressed genes, along with the numbers of significantly upregulated or downregulated genes for each group comparison between *ΔelyC* mutant and WT cells grown at 21°C versus 37°C across all six comparisons. Overlapping regions indicate genes uniquely expressed in different condition comparisons. Detailed data is in supplementary File S12. (H) A heatmap displays the most significant KEGG pathways and corresponding genes involved in amino and nucleotide sugar metabolism, showing upregulation solely in group condition (G1) 1-ΔelyC21/WT21. Detailed data is in supplementary File S13.

Our transcriptomic results demonstrate that various cell envelope modifications are affected at low temperatures in *E. coli*. The absence of ElyC evokes a highly defective cellular envelope when *ΔelyC* cells grow at 21°C. Our phenotypic studies reveal that for the first time, the ElyC factor is important for PG biosynthesis at 37°C as well. Additionally, in the both previous study and the current work we have shown that when the *ΔelyC* strain is grown at 21°C and 37°C, there is a significant accumulation of protein aggregation, compared to the wild-type strain. The absence of ElyC reduces the rate of PG biosynthesis at both temperatures with a more pronounced effect at the low temperatures, likely due to changes in the composition of inner membrane phospholipids. Thus, ElyC plays a crucial role in achieving optimal PG production rates at both high and low temperatures. While many mechanistic questions remain to be answered, our study provides compelling evidence that ElyC plays a significant role in *E. coli* cell envelope biogenesis and subsequent PG production through distinct mechanisms. This research opens new avenues for understanding how mutations in cell envelope genes affect expression levels in bacterial cell envelopes, particularly in the context of antibiotic resistance, and for developing effective therapeutic approaches.

## MATERIALS AND METHODS

### *E. coli* strains, culture media, and growth conditions

The strains used in this study include the *Escherichia coli* wild-type strain MG1655 (*rph1 ilvG rfb-50)* {Guyer, 1981 #12} along with its derivative *elyC::frt* (EM9) and *mrcB::frt* (MM39) {Paradis-Bleau, 2014 #10}. All cells were grown in Luria-Bertani (LB) medium containing 1% NaCl at either 21°C or 37°C, with continuous shaking at 250 rpm. Pre-cultures were initiated in LB medium and incubated at 37°C.

### Assessment of growth phenotypes and microscopic analysis of cell shape

To assess the growth of the Δ*elyC* mutant and wild-type *E. coli* strains in liquid media at 21°C and 37°C, single colonies of the mutant and WT were cultured in 6 ml LB in test tubes overnight at 37°C in an orbital shaking incubator. Subsequently, cells were diluted to an initial OD_600_ of 0.02 in LB medium and cultured in 25 ml of LB in 250 ml flasks at both 21°C and 37°C with continuous agitation at 250 rpm. The growth of cells was continuously monitored by measuring the OD_600_ at various time points over several hours following inoculation. At an OD_600_ of 0.5, just before the growth decline was observed for the Δ*elyC* mutant at 21°C, 3 µl of both mutant and WT cultures were spotted on 0.8% agarose pad. Images were captured using phase-contrast microscopy on a Nikon Eclipse E600 microscope equipped with a 100X oil immersion objective and a DS-Ri2 Nikon camera. Subsequent image analysis and cell width measurements were performed using Image J software (https://imagej.nih.gov/ij/). The average variation in cell width was determined for 100 cells of both WT and Δ*elyC* (at OD_600_ 0.5) at both 21°C and 37°C and recorded. Statistical data analysis was carried out based on three independent experiments, comparing the mutant to the WT cells.

### Peptidoglycan isolation and Ultra Performance Liquid Chromatography (UPLC) analysis

The wild-type and Δ*elyC* overnight cultures were diluted to an OD_600_ of 0.02 in 500 ml of LB with 1% NaCl and grown at 21°C and 37°C with constant shaking at 250 rpm. The cells were cultured until they reached an OD_600_ of ∼ 0.5 and were then harvested. PG isolation was performed using the boiling SDS method as described previously {Alvarez, 2016 #13; Desmarais, 2013 #14}. In brief, the samples were boiled in 5% SDS for 2 hours, and the sacculi were washed repeatedly with MilliQ water by ultracentrifugation (110,000 rpm, 10 min, 20°C). Subsequently, the isolated sacculi were treated with pronase E (100 μg/ml) to eliminate Braun’s lipoprotein. After adding SDS to a final concentration of 1% (w/v), the mixtures were heat-inactivated, and detergent was removed through additional washing steps as described. The samples were then treated with muramidase (100 μg/ml) for 16 hours at 37°C. The muramidase digestion was stopped by boiling, and coagulated proteins were removed by centrifugation (10 min, 14,000 rpm). The supernatants were initially adjusted to pH 8.5-9.0 using a sodium borate buffer, followed by the addition of sodium borohydride to a final concentration of 10 mg/ml. After a 30-minute reduction at room temperature, the pH of the samples was adjusted to 3.5 using orthophosphoric acid.

For UPLC analysis of muropeptides, a Waters UPLC system (Waters Corporation, USA) was utilized, equipped with an ACQUITY UPLC BEH C18 Column, 130Å, 1.7 μm, 2.1 mm **×** 150 mm (Waters, USA), and a dual-wavelength absorbance detector. Muropeptides were detected at 204 nm. Muropeptides were separated at 35°C using a linear gradient from buffer A (phosphate buffer 50 mM, pH 4.35) to buffer B (phosphate buffer 50 mM, pH 4.95 methanol 15% v/v) over a 20-minute run, with a flow rate of 0.25 ml/min. Relative PG amounts were calculated by comparing the total chromatogram intensities (total area) from three biological replicas, which were normalized to the same OD_600_ and extracted in the same volumes, as there is a direct correlation between muropeptide content and detected intensity. Muropeptides were identified by comparison with standard chromatograms of known composition, and their relative abundances were calculated as the relative area of the corresponding peak compared to the total area of the normalized chromatogram. The injection volume was optimized to ensure the signal fell within the system’s detection limits.

The PG chromatographic profiles were visually identical for all samples (Fig. 2B). Statistical analysis using the Student’s t-test revealed no significant differences in muropeptide composition and relative abundances between the Δ*elyC* and the wild-type cells, regardless of the different temperature growth conditions (21°C and 37°C). The relative abundance of the muropeptide peaks of interest is provided in Table S1. The percentage of cross-linking in each sample was consistent, approximately ∼30%, and a similar abundance obtained from the muropeptide fraction terminating with 1,6-anhydro, indicating that the average glycan strand length was comparable between the Δ*elyC* mutant and WT strains at both tested temperatures (Fig. 2C). The abundance of muropeptides containing the outer membrane Braun’s Lipoprotein (Lpp) moiety was consistent across all samples.

### Determination of β-lactam antibiotics sensitivity

The minimum inhibitory concentration (MIC) values of various β-lactam antibiotics against wild-type MG1655 and mutant strains, EM9 (Δ*elyC*) and MM39 (Δ*mrcB*), were assessed in LB 1% NaCl liquid media supplemented with different concentrations of antibiotics as specified. The cultures were initiated with a low bacterial concentration (2 x 10^6^ CFU/ml) and incubated at 37°C for 18 hours, after which growth was evaluated. To further validate increased sensitivity to β-lactam antibiotics, 30 ug/ml of ampicillin or cephalexin was introduced during the exponential growth phase of the wild-type MG1655, EM9 (Δ*elyC*), and MM39 (Δ*mrcB*) strains at 37°C.

### Assessment of protein aggregation

Aggregated proteins were isolated following the procedure outlined in {Tomoyasu, 2001 #78}, with a modification as previously described {Kouidmi, 2018 #67}. Wild-type MG1655 and Δ*elyC* were initially cultured overnight in LB at 37°C. Subsequently, they were inoculated into 25 ml fresh LB media in the morning, targeting an OD_600_ of 0.02, and grown at 37°C until reaching an OD_600_ between 0.3 and 0.35. The cells were then transferred to a pre-cooled 50 ml conical tube and centrifuged at 6000 g for 10 minutes at 4°C. The resulting pellets were resuspended in 40 μl of buffer A (comprising 10 mM potassium phosphate buffer [pH 6.5], 1 mM EDTA, 20% [wt/vol] sucrose, 1 mg/ml of lysozyme, and 1.4 mM phenylmethylsulfonyl fluoride [PMSF]), followed by 30-minutes of incubation on ice. Subsequently, 360 μl of buffer B (containing 10 mM potassium phosphate buffer [pH 6.5], 1 mM EDTA, and 1.4 mM PMSF), was added to the cells, mixed, and sonicated while being cooling using a Sonics Materials VCX 400 sonifier (microtip, level 2, 50% duty, 10 cycles). Intact cells were separated by centrifugation at 2,000 g for 15 minutes at 4°C. The insoluble fraction, encompassing membrane and aggregated proteins was obtained by centrifugation at 15,000 g for 20 minutes at 4°C. The pellet fractions were frozen, resuspended in 1 ml of buffer B via brief sonication, and then centrifuged at 15,000 g for 20 minutes at 4°C. This washing procedure was repeated once more. Then, 200 μl of 10% (wt/vol) NP-40 was added, and the mixture was incubated for 4 hours at 4°C with agitation. The aggregated proteins were then isolated by centrifugation at 15,000 g for 30 minutes at 4°C. The soluble fractions, containing membrane proteins, were conserved, while the insoluble pellets (containing aggregates) were resuspended in 200 μl of buffer B. Protein quantification was performed using the Bradford method {Bradford, 1976 #79}, employing the BIO BASIC INC. Kit #SK3041, Version 3.0.

### Transcriptomic analysis by RNA-seq

In triplicate cultures, both the wild-type and Δ*elyC* mutant strains were grown in 250 ml of LB 1% NaCl liquid media at 250 rpm, maintaining temperatures at 21°C and 37°C until reaching the early-log phase (OD_600_ of 0.30 to 0.35). The samples were immediately centrifuged (6000 g) at 4 °C for 5 minutes. The pellets were washed once with cold TES buffer (comprising 100 mM NaCl, 1 mM EDTA, 10 mM Tris hydrochloride [pH 7.4]), followed by centrifugation. The cells were then snap-frozen in liquid nitrogen and stored at −85 °C. Total RNA was purified using the RNeasy Mini Kit (QIAGEN) with the RNA protect Bacteria reagents (QIAGEN), following the manufacturer’s protocol. To eliminate any residual DNA, the RNA samples underwent treatment with the RNase-Free DNase set from QIAGEN. The purified RNA was ultimately resuspended in RNase-Free Water (QIAGEN), and its concentration was determined at 260 nm using a NanoDrop ND-1000 spectrophotometer. The RNA Ribosome Integrity Numbers (RIN) and purity of the RNA samples were assessed using the Agilent Bioanalyzer 2100 and RNA 6000 Pico Kit. All twelve samples exhibited RIN numbers ranging from 9.3 to 10. To remove ribosomal RNA (rRNA), the Ribo-Zero Magnetic Gold Kit for Gram-negative Bacteria (Illumina) was employed. RNA-seq libraries were prepared using the KAPA RNA stranded Kit (Roche-Nimblegen), and their quality and quantity were assessed with a BioAnalyzer using the High Sensitivity DNA Kit (Agilent). Sequencing was performed using an Illumina HiSeq2000 machine, in a 75-nucleotide paired-end mode, at the Bioinformatics Core Facility, IRCM, University of Montreal, Canada. Trimming of Poor-quality reads was carried out using Trimmomatic {Lohse, 2012 #15}, and the quality of the reads, before and after trimming, was assessed with FASTQC version 0. 11. 5. The reads were aligned to the *E. coli* sub-strain MG1655 reference genome (GCA000801205.1) using Bowtie2 version 2. 2. 9 {Langmead, 2012 #16}. The reference genome for alignment, and the GTF annotation file for read counting were obtained from Ensembl (release 27). The aligned data were saved as BAM files. The raw read counts for each gene were calculated using *featureCounts* 1.5.1 {Liao, 2014 #45}{Langmead, 2012 #34}. Subsequently, normalization and determination of differential gene expression (fold change /Log_2_ fold change) were performed, along with *p*-value (< 0.05) calculations, using DESeq 2. 1. 14.0 in the R program {Anders, 2010 #29;Love, 2014 #46;Love, 2014 #46}{Liao, 2014 #45}. The output from DESeq2 included the raw counts normalized relative to the total number of reads. Differentially expressed genes were determined based on *p*-values (< 0.05), and as part of multiple comparison testing, significant Benjamini-Hochberg adjusted *p*-values (p-adj) < 0.05 (5% false discovery rate) were considered. Additionally, the counts were normalized relative to library size, and genome-wide occupancy was quantified in density units of fragments per kilobase of exon model per million reads mapped (FPKM). To assess gene expression, we utilized BV-BRC bioinformatics resource (Olson et al. 2023; https://www.bv-brc.org/). Boxplot (Spitzer et al. 2014; http://shiny.chemgrid.org/boxplotr/), GraphPad Prism 10, and R software were used to generate dendrogram clustering, MA plots, scatter plot, PCA, biplot, Pearson correlation, volcano plots, and heat maps. The DAVID bioinformatics platform (Sherman et al. 2022; https://david.ncifcrf.gov/), was employed for gene annotation, functional clustering, enrichment, and pathway analysis.

### Statistical analysis

Statistical analyses for cell width and peptidoglycan data were conducted using GraphPad PRISM^®^ Software (Inc., San Diego CA, www.graphpad.com). The significance of the results was determined through unpaired Student’s t-tests, and this analysis was confirmed with three independent replicates to ensure the reproducibility of the result. Transcriptomics data analysis and the generation of graphs were performed using DESeq 2. 1. 14.0 in the R program following established methodologies {Love, 2014 #46;Anders, 2010 #29}{Anders, 2010 #29;Love, 2014 #46;Love, 2014 #46}, and GraphPad PRISM^®^ Software version 10.

### Data availability

The raw RNA-Seq data in FASTQ format and gene counts from this study have been deposited to the NCBI Gene Expression Omnibus (GEO) under GEO accession number GSE141694.

## SUPPLEMENTAL MATERIAL

Supplemental material for the paper is available at the following link: https://

**SUPPLEMENTAL FILE 1,** PDF file, Size: XXXXX MB.

All study data, including Source Data for Figures 1-7 (XLS and Word formats) and supplementary figures S1-S8, are included in the manuscript and supporting files.

**FIG 7.**
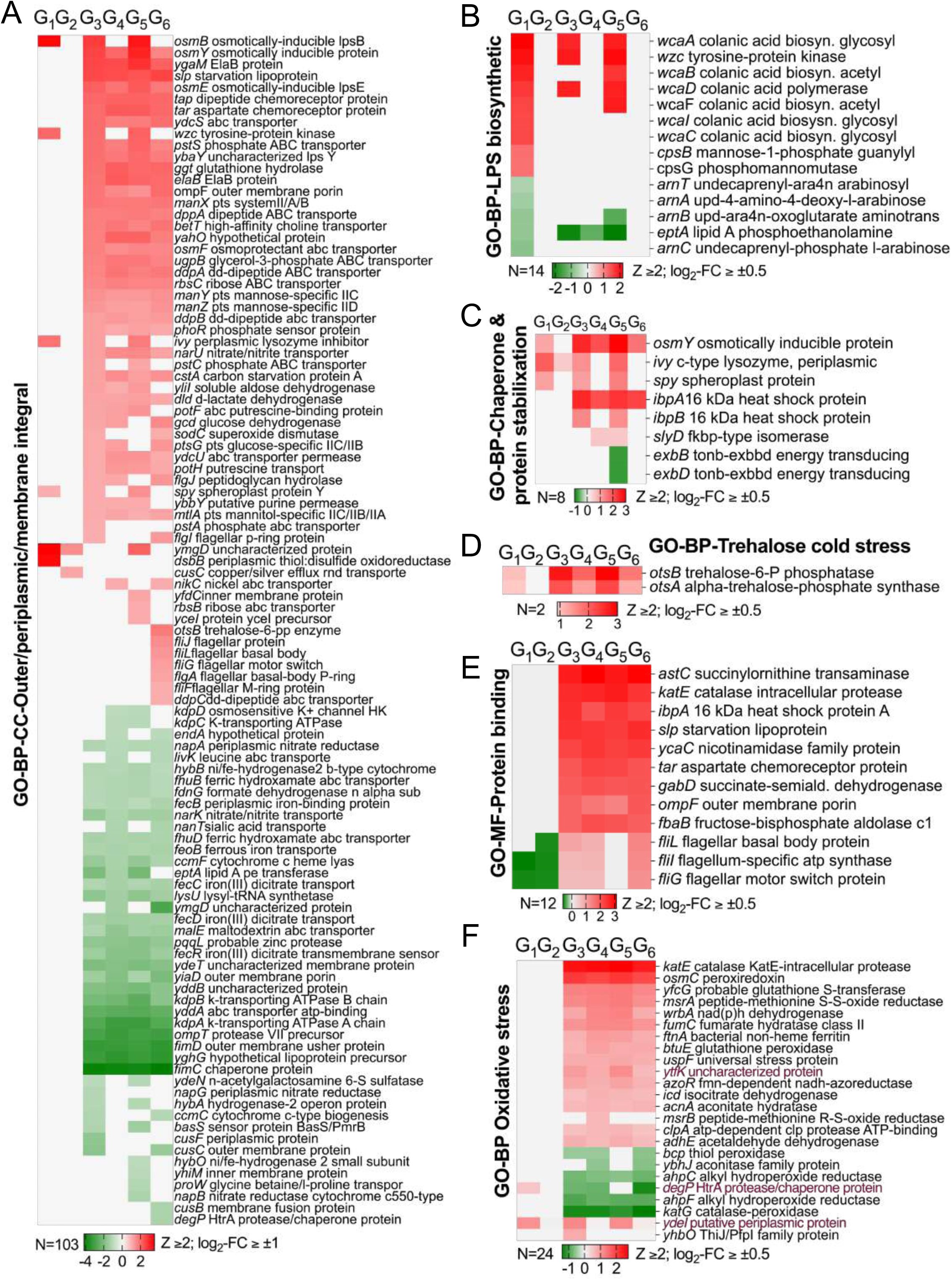
Illustrates how the *ΔelyC* mutant and growth temperature shift impact the transcriptional expression of mRNAs encoding key molecular functions, biological processes, and cellular components involved in cell envelope biogenesis, and environmental stress. (A-F) Heatmaps highlight up/downregulated genes in outer membrane, periplasmic, and membrane integrity enriched in GO biological processes (BP) and cellular components (CC), lipopolysaccharide biosynthesis (B), chaperone/protein stabilization (C), trehalose biosynthesis under cold stress (D), protein binding (E), oxidative stress (F), enriched in GO molecular function and biological process (see FIG 5-B-D, FIG 6-B, and FIG S5-7), defining functional categories across six groups (G1-G6). Values show log_2_-fold change (log2-FC) ≥ ±0.5 or ≥ ±1 and Z-score (Z) ≥ 2, using color scales: green for down-regulated, red for up-regulated, silver for no significant change.

## ACKNOWLEDGMENTS

We would like to thank Dr. Marc Drolet, Dr. Elitza Tocheva, and Rim Marrakchi for their feedback on the first draft of this manuscript. We also extend our thanks to Irene Asensio Gudina for her assistance with the microscopic images and the CMI for providing β-lactam antibiotics. We thank Negar Baizapour for her invaluable help in data mining and managing large datasets, enabling comprehensive analysis and exploration. We appreciate Clark Cuccinell at BVBRC for ensuring data matrix compatibility with the differential expression service. The research conducted in the Paradis-Bleau lab was funded by the Natural Sciences and Engineering Research Council of Canada (NSERC) and the Université de Montréal. Research in the Cava lab was supported by the Knut and Alice Wallenberg Foundation (KAW), the Laboratory of Molecular Infection Medicine Sweden (MIMS), the Swedish Research Council, and the Kempe Foundation.

The funders had no role in study design, data collection and interpretation, or the decision to submit the work for publication.

## SUPPLEMENTARY FIGURE LEGENDS

**FIG S1.**
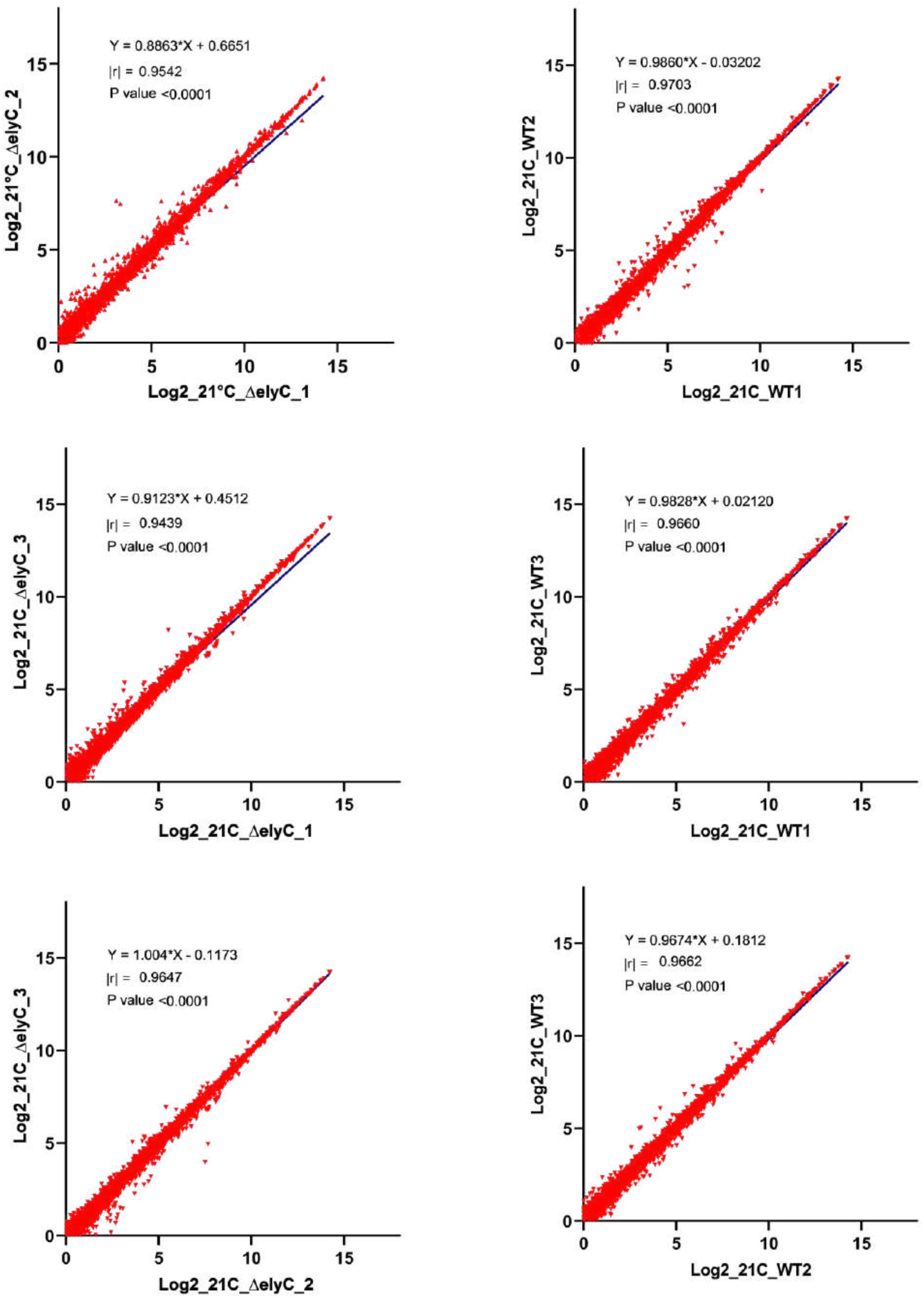
(A, B, C, D, E, F) Scatter plots demonstrate robust correlation coefficients (|r| > 0.94 to 0.98, p <0.0001) between biological replicates of Δ*elyC* mutant and WT cells grown at 21°C.

**FIG S2.**
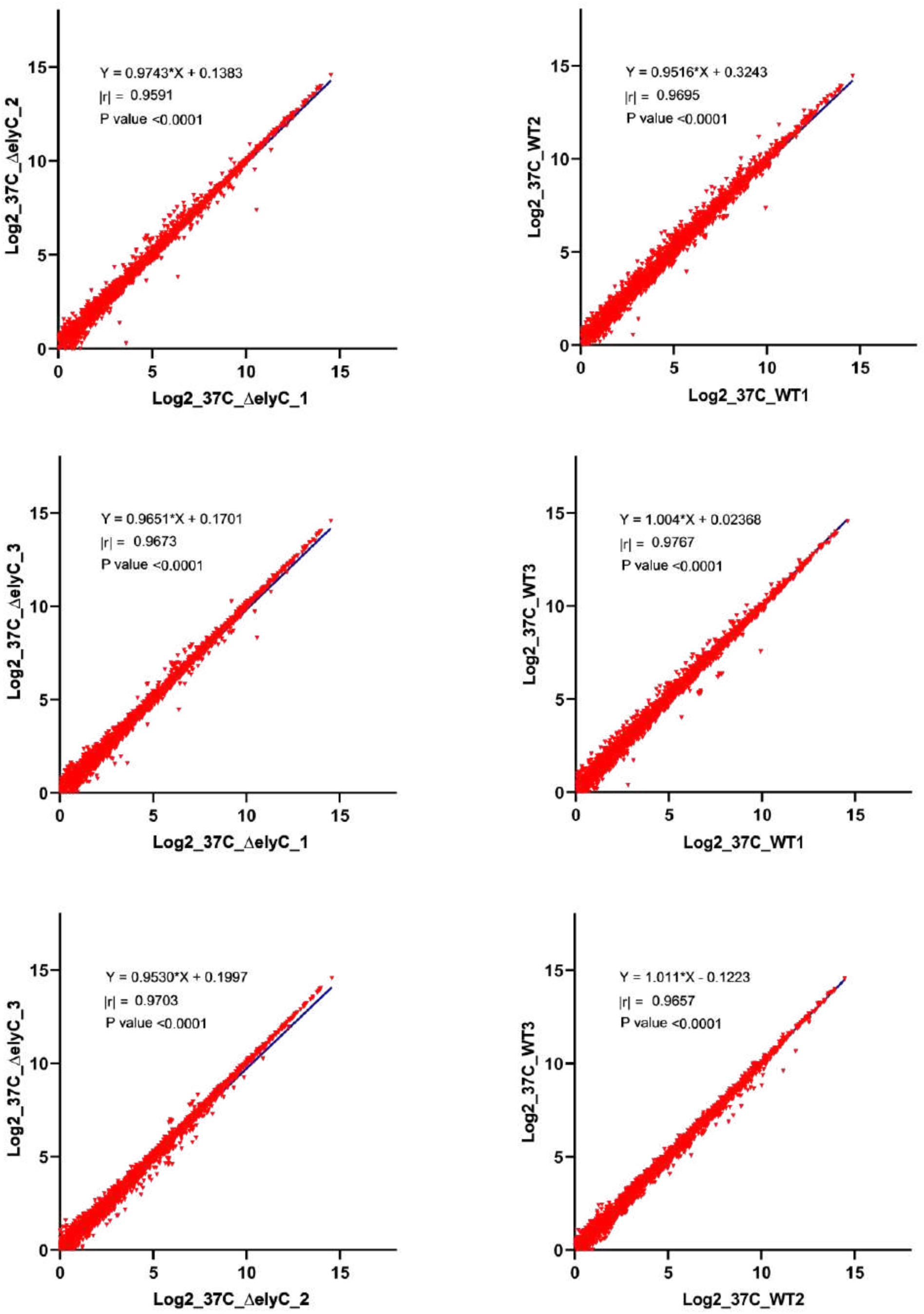
(A, B, C, D, E, F) Scatter plots reveal robust correlation coefficients (|r| > 0.94 to 0.98, p <0.0001) between biological replicates of Δ*elyC* mutant and WT cells grown at 37°C.

**FIG S3.**
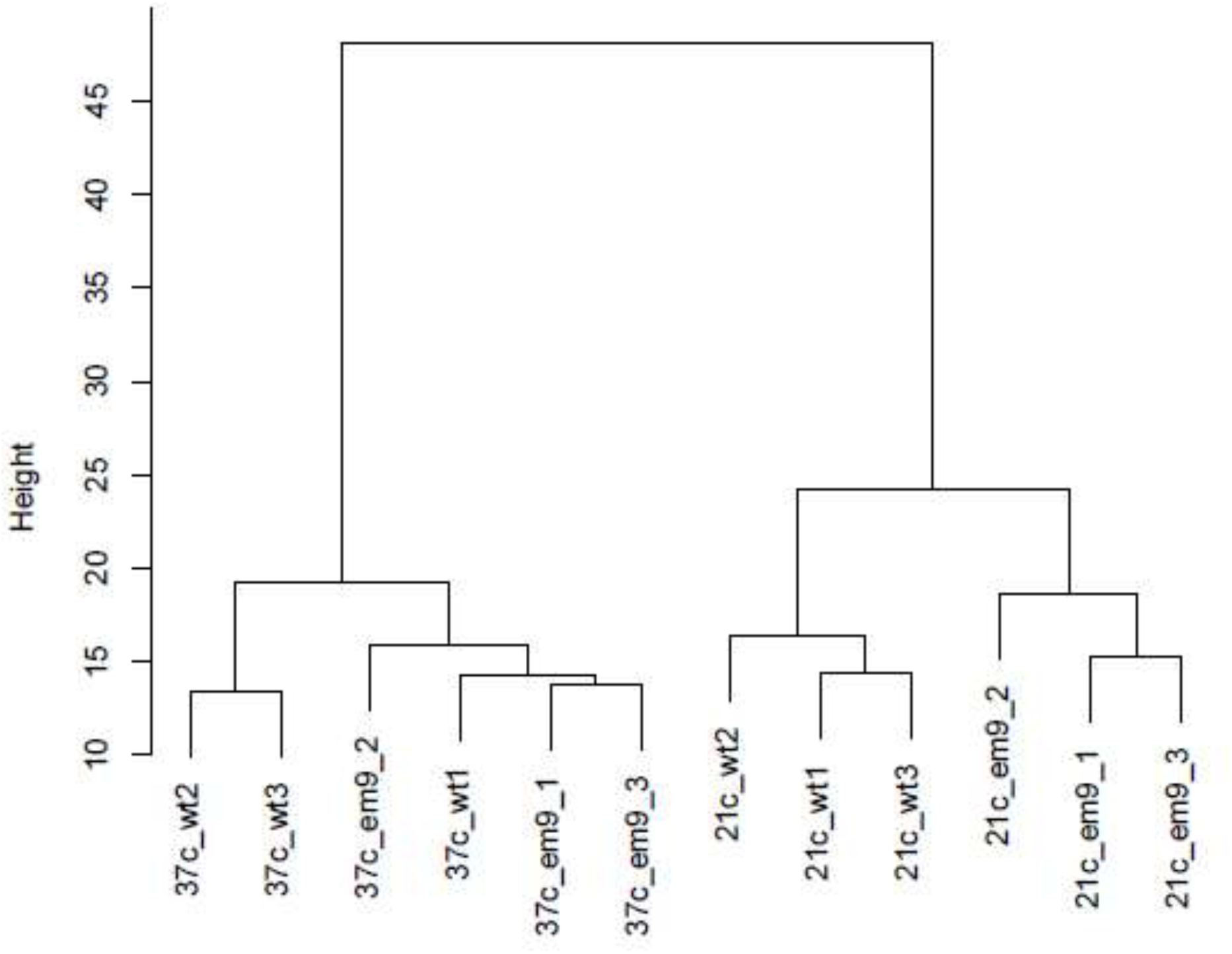
Hierarchical clustering of samples based on gene counts after variance stabilization transformation with DESeq2, illustrating the relationship between different experimental conditions.

**FIG S4.**
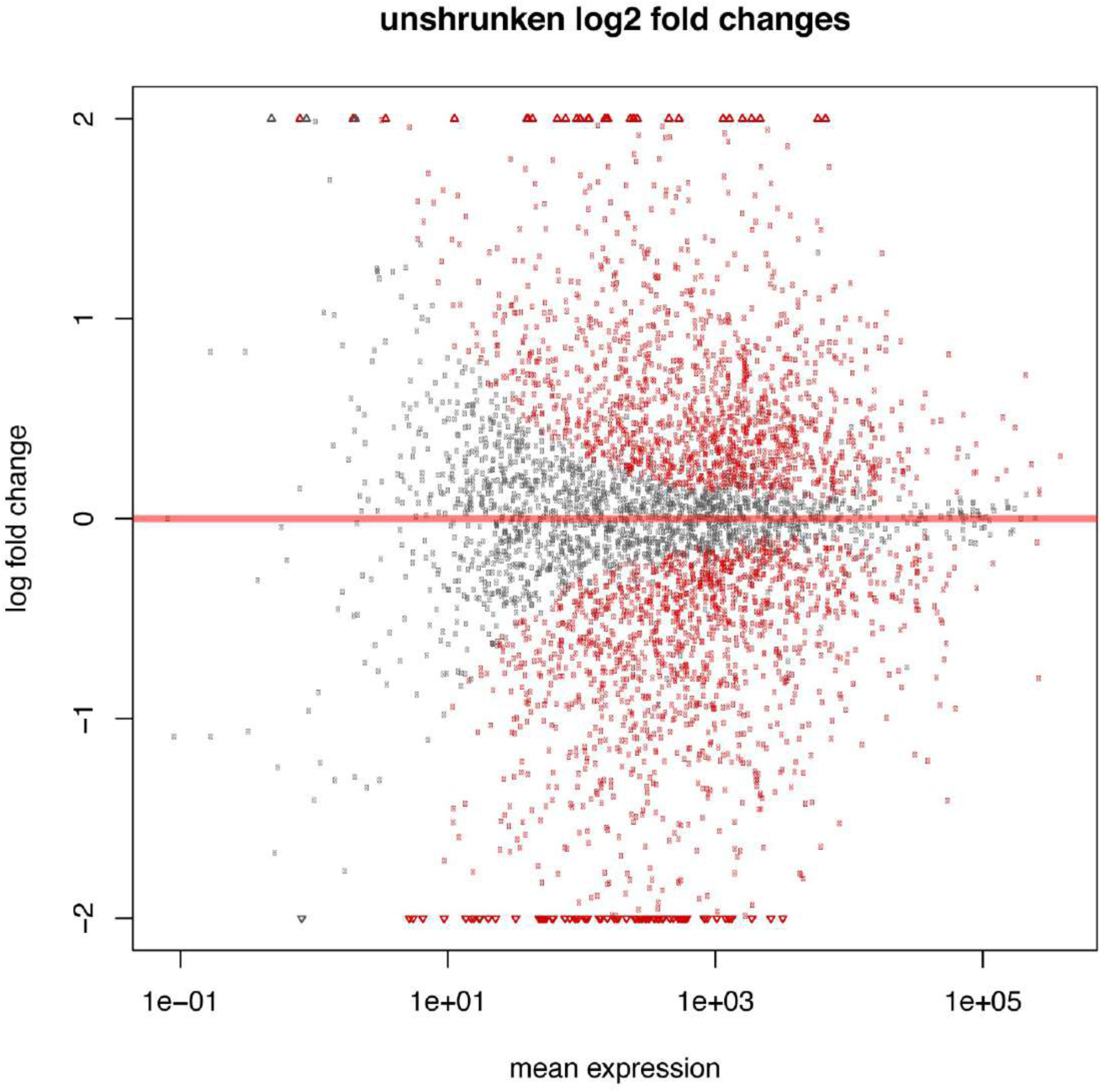
MA plot generated using the R package, depicting the distribution of log_2_-fold change in overall gene expression based on mean normalized counts.

**FIG 6-D FIG supplement S5-A-B.**
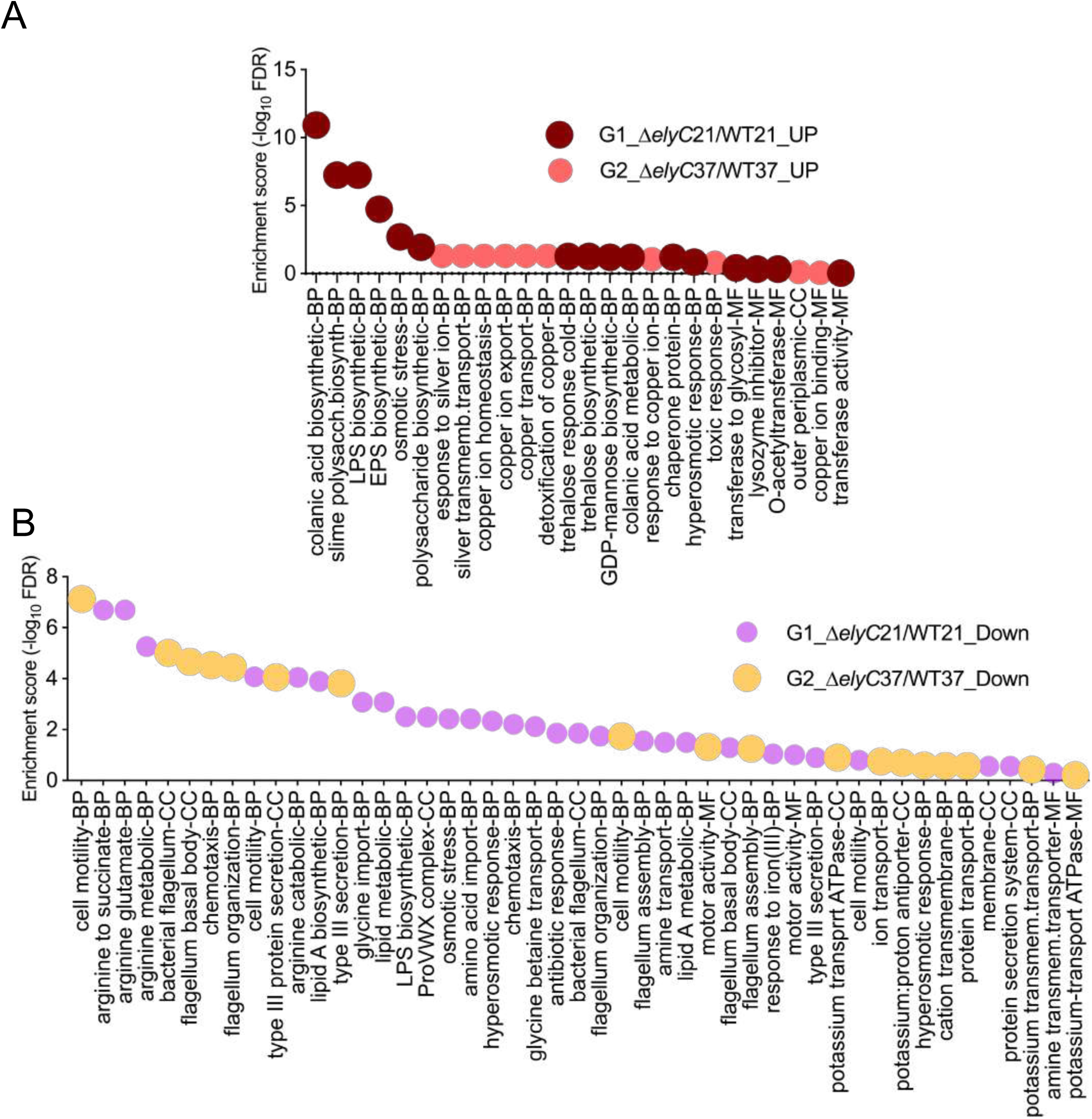
displays gene ontology (GO) enrichment analysis in the subset of 172 differentially expressed mRNAs (FDR <0.316; Z ≥2; log2(FC) ≥ ±0.5) induced by the *ΔelyC* mutant and WT strain at 21^∘^C and 37^∘^C. Bubble plots (A-B) depict enriched GO terms in biological processes (BP), cellular components (CC), and molecular functions (MF) for upregulated genes in group G1-*ΔelyC*21/WT21 and group G2-*Δely*C37/WT37 (A), and downregulated genes in the same groups (B). The y-axis shows the negative log10 of adjusted p-values (FDR) for GO enrichments, while the x-axis lists the GO terms. Additional data are available in the XLS files 8 and 9_S5-S7.

**FIG 6-E FIG supplement S6-C-D.**
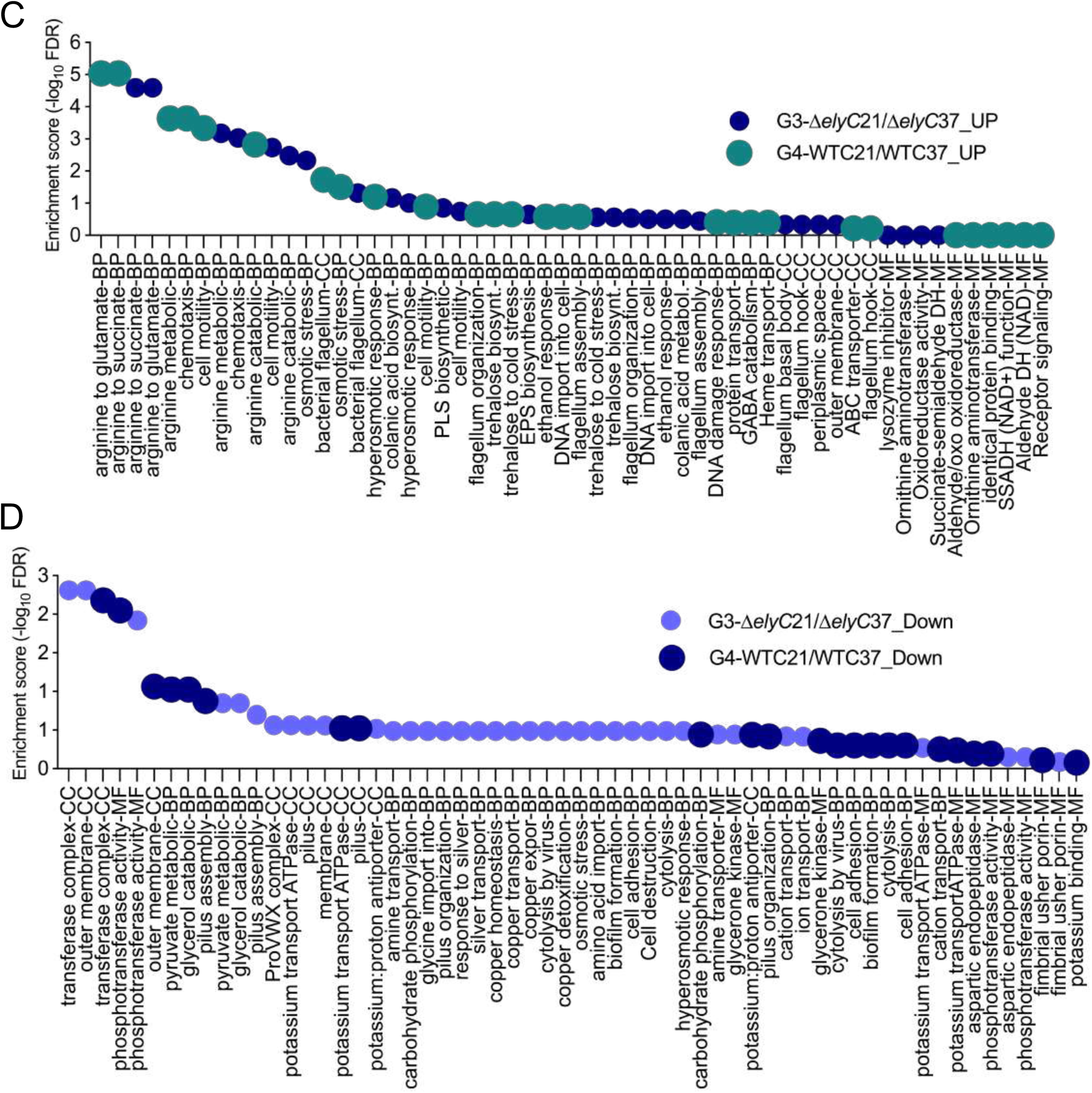
displays gene ontology (GO) enrichment analysis in the subset of 172 differentially expressed mRNAs (FDR <0.001; Z ≥2; log2(FC) ≥ ±0.5) induced by the *ΔelyC* mutant and WT strain at 21^∘^C and 37^∘^C. Bubble plots (C-D) depict enriched GO terms in biological processes (BP), cellular components (CC), and molecular functions (MF) for upregulated genes in group 3-*ΔelyC*21/*ΔelyC*37 and group 4-WTC21/WTC37 (C), and downregulated genes in the same groups (D). The y-axis shows the negative log10 of adjusted p-values (FDR) for GO enrichments, while the x-axis lists the GO terms. Additional data are available in the XLS files 8 and 9_S5-S7.

**FIG 6-F FIG supplement S7-E-F.**
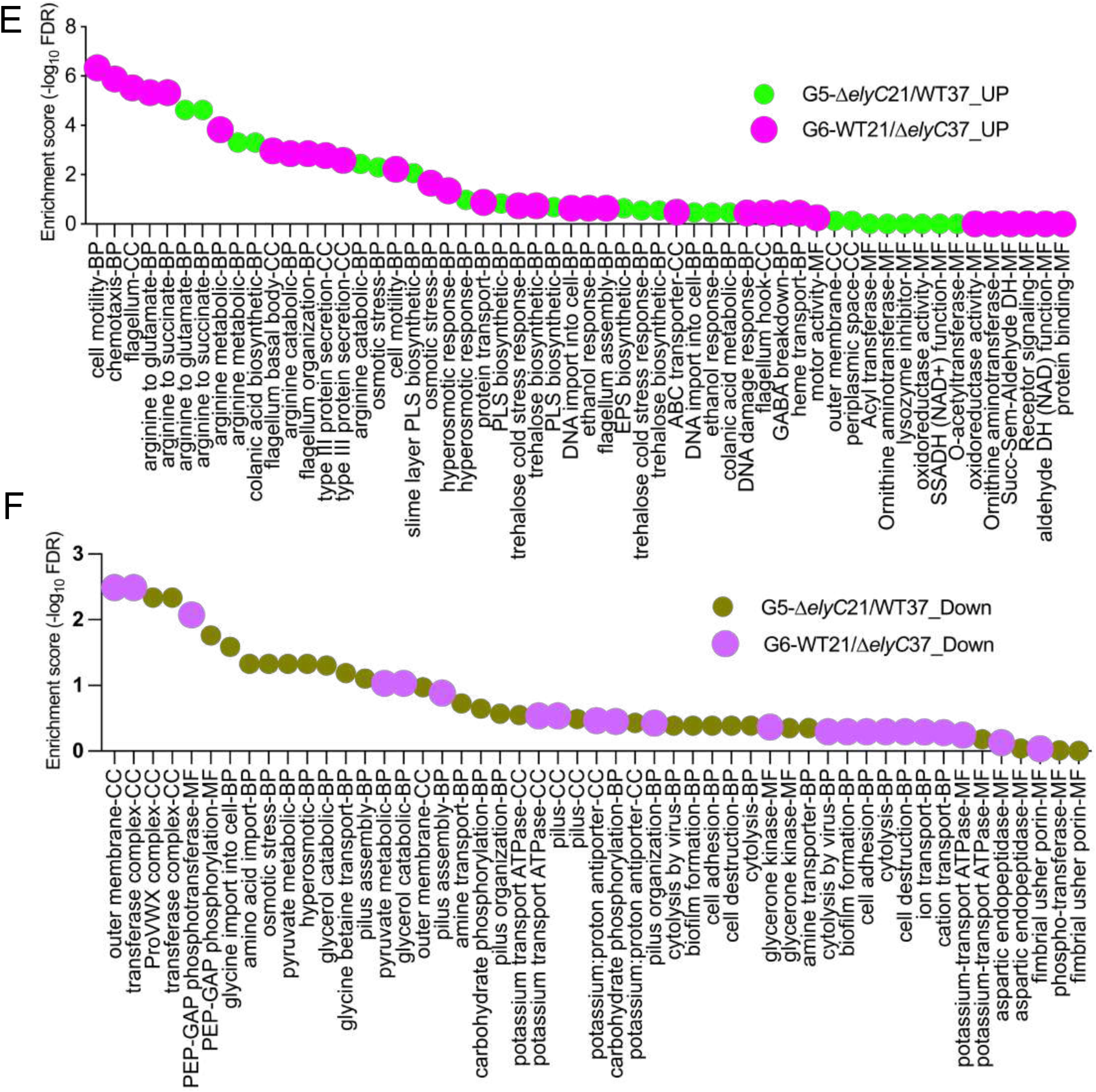
displays gene ontology (GO) enrichment analysis in the subset of 172 differentially expressed mRNAs (FDR <0.001; Z ≥2; log2(FC) ≥ ±0.5) induced by the *ΔelyC* mutant and WT strain at 21^∘^C and 37^∘^C. Bubble plots (E-F) depict enriched GO terms in biological processes (BP), cellular components (CC), and molecular functions (MF) for upregulated genes in group 5-*ΔelyC*21/WT37 and group 6-WT21/*ΔelyC*37 (E), and downregulated genes in the same groups (F). The y-axis shows the negative log10 of adjusted p-values (FDR) for GO enrichments, while the x-axis lists the GO terms. Additional data are available in the XLS files 8 and 9_S5-S7.

**FIG S8-A-E.**
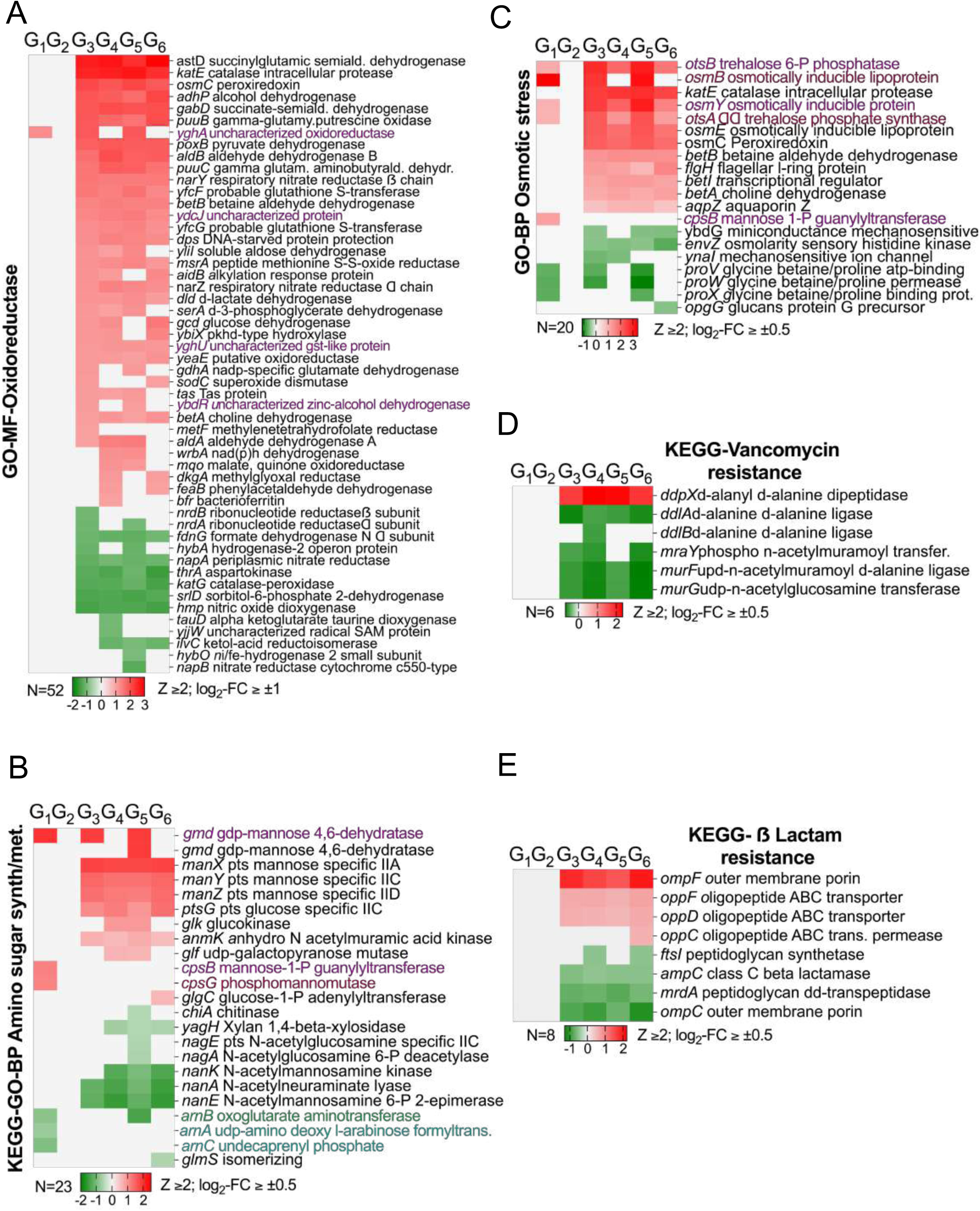
Display how the *ΔelyC* mutant and growth temperature shift impact the transcriptional expression of mRNAs encoding key molecular functions, biological processes, and cellular components indirectly involved in cell envelope biogenesis and antibiotic resistance stress. (A-E) Heatmaps highlight up/downregulated genes in oxidoreductase (A), amino sugar metabolism/nucleotide sugar synthesis (B), osmotic stress (C), vancomycin resistance (D), and beta-lactam resistance (E), enriched in GO biological processes (BP), molecular function (MF), KEGG pathways, defining functional categories across six groups (G1-G6) (see FIG 5-B-D, 6-C-F, and FIG S5-7). Values represent log2-fold change (log2-FC) ≥ ±0.5 or ≥ ±1 and Z-score (Z) ≥ 2, using color scales: green for down-regulated, red for up-regulated, silver for no significant change.

## TABLES SOURCE DATA

**Additional files 1-2**

**Table S1** Muropeptide relative abundance.

Note: The relative molar abundance of the depicted peaks in the chromatograms (Fig. 2E) is calculated based on the relative area of each peak and expressed as the mean percentage value from three independent cultures. Statistical analysis was conducted using Student’s t-test, comparing each sample to the WT in each condition.

**Table S2** Raw read statistics of the RNA-seq samples.

**Additional file S3**_1866_1710_172_Volcano_diffexp_gene_mtx

**Additional file 4**_FIG 5_B-C-D_GO_Functional Annotation Chart

**Additional file 5**_FIG 5_E_F_KEGG_Pathways_All 1710

**Additional file 6**_FIG 6_A

**Additional file 7**_FiIG 6_B_GO_172 genes

**Additional file 8**_FIG 6_S5-S7_Bubble_G1_6_UP

**Additional file 9**_FIG 6 S5-S7_Bubble_G1_6_Down

**Additional file 10**_FIG 6_C_KEGG_All_G_ 172 genes_Z_2_FC_ ≥ ±0.5.

**Additional file 11**_FIG 6_D-E-F_KEGG Up_Down_6 groups_172 genes

**Additional file 12**_FIG 6_G_Source Data

**Additional file 13**_FIG 6_H_Heatmap 172 genes z_2_FC_≥ ±0.5 PE pathways

